# Brain endothelial PRMT5-ANGPTL4 axis regulates cerebellar inhibitory synaptogenesis and motor coordination

**DOI:** 10.1101/2025.11.18.688985

**Authors:** Jinjin Lu, Yizhe Zhang, Yijie Hao, Huimin Ning, Zizhao Fan, Shiqiang Hao, Yaxiong Cui, Yuying Han, Yunting Cai, Wenjia Liu, Haiou Ao, Shunshun Xiu, Jun Wang, Xiao Yang

## Abstract

Brain vasculature is essential for central nervous system function, but its specific roles in synaptic development and motor function regulation are poorly understood. In this study, we identify a cerebrovascular signaling axis wherein endothelial PRMT5 critically regulates inhibitory synaptogenesis and motor coordination. Cerebrovascular-specific deletion of *Prmt5* gene leads to excessive inhibitory synaptic input onto Purkinje cells and progressive motor deficits in mice. Mechanistically, PRMT5 deficiency epigenetically upregulates the expression of the secreted factor ANGPTL4 through reduced symmetric dimethylation of H3R8 and H4R3, along with increased H3K9 acetylation at the *Angptl4* promoter, thereby driving excessive inhibitory synaptogenesis onto Purkinje cells. Importantly, *Angptl4* deletion in brain endothelium normalizes inhibitory synaptic input onto Purkinje cells and restores motor coordination in PRMT5-deficient mice. Together, these findings define the endothelial PRMT5-ANGPTL4 axis as a key regulator of cerebellar inhibitory circuitry and motor function, highlighting cerebrovascular mechanisms as potential therapeutic targets for motor coordination disorders.

Teaser

Cerebrovascular PRMT5 prevents excessive inhibitory synaptogenesis via epigenetic repression of *Angptl4*.

## Introduction

Brain vasculature plays essential roles in central nervous system (CNS) development and homeostasis, serving not only as a conduit for oxygen/nutrient delivery and waste removal but also as an active signaling hub within neurovascular units (NVUs) (*1, 2*). As integral components of the NVU, cerebrovascular endothelial cells (ECs) actively orchestrate key neurodevelopmental processes, including neuronal proliferation, maturation, spatial positioning, circuit assembly, and synaptic plasticity (*3–6*). These effects are mediated primarily through the secretion of paracrine signaling molecules that act directly on neural cells (*4–8*) or through the dynamic regulation of substance transport across the blood-brain barrier (BBB) (*9–13*). Despite growing recognition of their regulatory roles, the specific contributions of ECs to cerebellar circuit function and motor coordination remain poorly understood.

The cerebellum, a pivotal hub for sensorimotor integration and coordination, depends on the precise synaptic organization of its cortical circuits (*14*). Purkinje cells (PCs), the sole output neurons of the cerebellar cortex, integrate excitatory inputs from climbing fibers (CFs) and parallel fibers (PFs) with finely tuned inhibitory inputs from molecular layer interneurons (MLIs) (*14–16*). Degeneration and dysfunction of PCs represent hallmark pathological features in a range of neurological disorders associated with motor impairments (*17–20*). Beyond cell-autonomous impairments, accumulating evidence indicates that circuit-level deficits upstream of PCs are sufficient to disrupt PC activity and motor output (*21–24*). Notably, a recent in vivo study demonstrates that excessive inhibitory synapses from MLIs onto PCs is sufficient to impair PC intrinsic excitability, trigger degeneration, and cause motor dysfunction (*21*), underscoring the critical importance of properly regulated inhibitory synaptogenesis for cerebellar function. While previous studies have identified multiple mechanisms underlying inhibitory synaptogenesis, including neuronal transmembrane adhesion molecules (*25–29*) and secreted factors (*30–32*), the known repertoire of these synaptic organizers within the brain microenvironment is primarily attributed to glial cells and neurons (*33*). The potential contribution of ECs remains largely unexplored. Given their proximity to neural circuits and their robust secretory capacity, ECs are poised to serve as potential sources of synaptogenic signals. Supportively, emerging evidence has identified EC-derived proteins as direct modulators of synaptic plasticity (*5, 8*). However, the specific role of ECs in cerebellar inhibitory synaptogenesis remains undefined.

Protein arginine methyltransferase 5 (PRMT5), a type II arginine methyltransferase that catalyzes the symmetric dimethylation of arginine residues (*34*), has been implicated in angiogenesis and vascular regeneration following acute ischemic injury (*35, 36*), yet its role in cerebrovascular signaling and neural circuit regulation is unknown. Here, we report a previously uncharacterized endothelial signaling pathway wherein PRMT5 critically constrains inhibitory synaptogenesis and maintains motor coordination. Using mice with cerebrovascular EC-specific deletion of *Prmt5*, we show that endothelial PRMT5 deficiency epigenetically upregulates the secreted factor ANGPTL4, which in turn drives excessive inhibitory synapse formation onto PCs and leads to progressive PC dysfunction and motor deficits. Importantly, brain endothelial-specific ablation of *Angptl4* normalizes inhibitory synaptic input onto PCs and rescues motor function in PRMT5-deficient mice.

## Results

### Endothelial PRMT5 deficiency induces progressive cerebrovascular abnormalities and motor deficits

To investigate the role of endothelial *Prmt5* in the brain, we generated *SP-A-Cre;Prmt5^flox/flox^* (*Prmt5^fl/fl^*) mutant mice and *SP-A-Cre;Prmt5^flox/+^* (*Prmt5^fl/+^*) control mice by crossing *Prmt5^flox/flox^* mice (*37*) with *SP-A-Cre* transgenic mice (*9, 38*). Cre recombinase activity was detected exclusively in CD31^+^ ECs, but not in NeuN^+^ neurons, GFAP^+^ astrocytes, or IBA1^+^ microglia in *SP-A-Cre;ROSA26^LSL-tdTomato^*(*SP-A-Cre^tdTomato^*) mice (fig. S1). Immunofluorescence staining confirmed the absence of PRMT5 in brain ECs of *Prmt5^fl/fl^*mice (fig. S2A), which was further verified at the mRNA level by qPCR (fig. S2B) and at the protein level by Western blot (fig. S2C) in isolated brain ECs.

We first examined whether *Prmt5* deletion in ECs affected cerebrovascular architecture by assessing vascular density and diameter through immunofluorescence analysis. Although *SP-A-Cre* expression begins at the embryonic stage (*39*), no significant cerebrovascular morphological abnormalities were observed in *Prmt5^fl/fl^*mice at postnatal day 20 (P20) (fig. S3, A to C). However, by P60, *Prmt5^fl/fl^*mice exhibited a modest but statistically significant increase in cerebrovascular diameter compared with controls (Fig. 1, A to C). Furthermore, by 3 months of age, *Prmt5^fl/fl^* mice displayed cerebrovascular malformations, predominantly in the cerebellar and thalamic regions but not the cortex, suggesting region-specific effects of PRMT5 deficiency on cerebrovascular structure (fig. S3D). These results demonstrate that cerebrovascular specific deletion of *Prmt5* leads to progressive cerebrovascular anomalies, underscoring its essential role in maintaining vascular homeostasis rather than in early cerebrovascular development.

**Figure 1.**
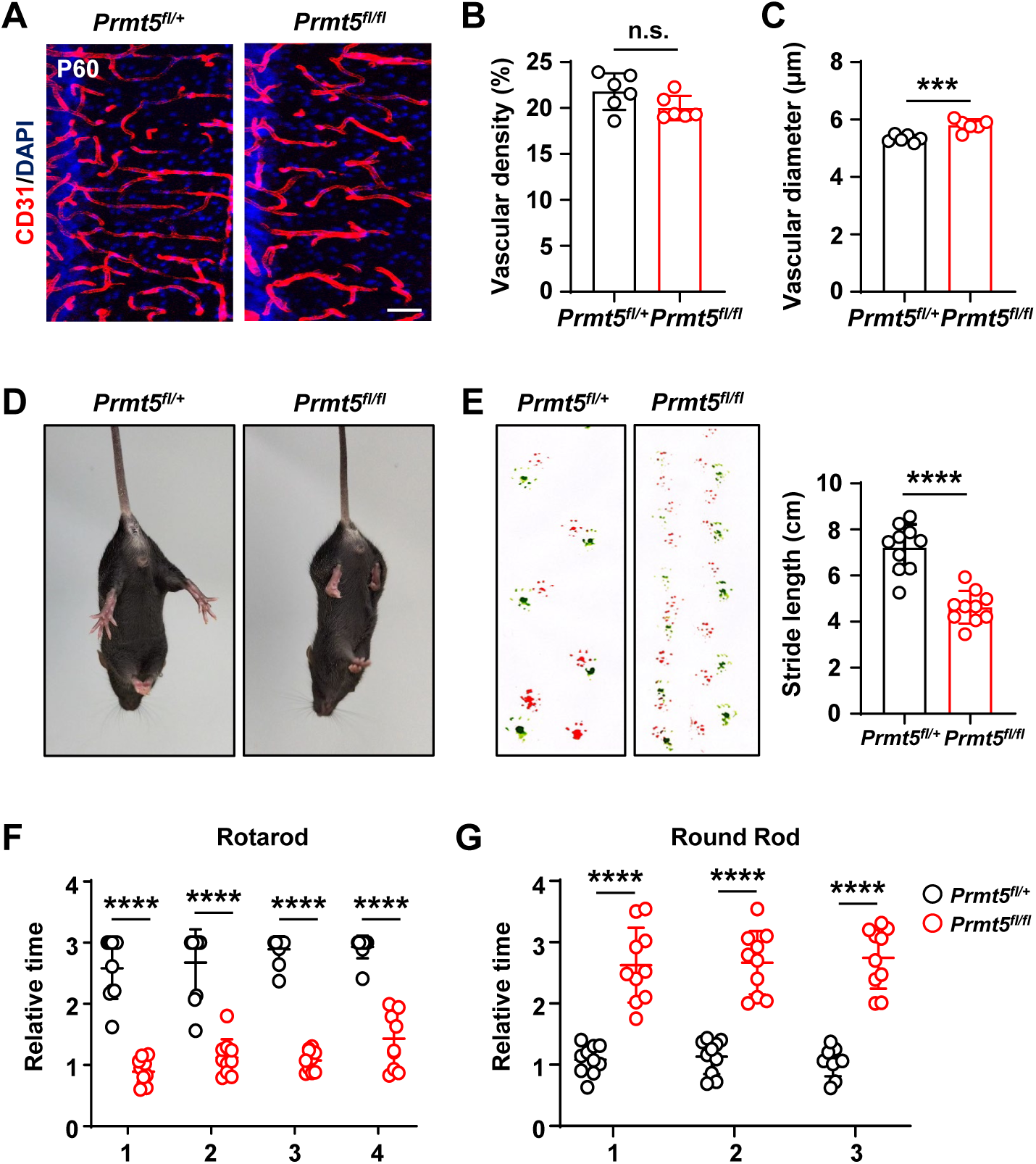
Loss of endothelial *Prmt5* leads to abnormal cerebrovascular structure and motor deficits. (A) Confocal images of CD31 (red) immunostaining showing cerebrovascular morphology in P60 *Prmt5_fl/+_* and *Prmt5_fl/fl_* mice. Scale bar, 50 µm. (B and C) Bar graphs showing cerebrovascular density (B) and diameter (C) in P60 *Prmt5_fl/+_* and *Prmt5_fl/fl_* mice. ***P < 0.001, n.s., not significant (mean ± SEM, n = 6 mice per group). (D) *Prmt5_fl/fl_* mice showed a limb clasping reflex in tail suspension test. (E) Representative footprints and quantification of stride length in P60 *Prmt5_fl/fl_* and *Prmt5_fl/+_* mice. Red, forepaws; green, hindpaws. ****P < 0.0001 (mean ± SEM, n = 10 mice per group). (F) Quantification of latency time spent on the rotarod. Data are presented from 4 independent trials. ****P < 0.0001 (mean ± SEM, n = 10 mice per group). (G) Quantification of time spent crossing the balance beam. Data are presented from 3 independent trials. ****P < 0.0001 (mean ± SEM, n = 10 mice per group).

Notably, *Prmt5^fl/fl^* mice exhibited a progressively exacerbating motor deficits, characterized by a wobbly posture and circular walking patterns at P60, ultimately culminating in an inability to maintain an upright stance on their hind limbs by 5 months of age (Video S1). Tail suspension revealed a limb clasping reflex in *Prmt5^fl/fl^* mice at P60 (Fig. 1D), a typical symptom of neurodegenerative defect (*40*). To quantify these motor deficits, we conducted a series of behavioral tests on male *Prmt5^fl/fl^* mice and control mice at P60. Footprint analysis showed significantly shorter stride length in *Prmt5^fl/fl^* mice compared to controls (Fig. 1E), consistent with ataxic locomotion. In the rotarod test, *Prmt5^fl/fl^*mice exhibited a significant reduction in latency to fall (Fig. 1F). Similarly, in the balance beam test, *Prmt5^fl/fl^* mice took longer time to cross the 11-mm-wide beam (Fig. 1G). Notably, these motor deficits were detectable as early as P20 (fig. S4) and progressively worsened with age.

Taken together, these findings demonstrate that endothelial-specific *Prmt5* deletion induces progressive cerebrovascular abnormalities and motor deficits, with detectable motor deficits emerging prior to vascular alterations. These results highlight that endothelial PRMT5 deficiency impairs motor coordination through abnormal neurovascular communication rather than direct vascular alterations or their secondary consequences.

### Increased inhibitory synapses on PCs in *Prmt5^fl/fl^* mice

Since the cerebellum is a central hub for sensorimotor coordination and was particularly vulnerable to *Prmt5* deficiency, we examined cerebellar morphology of *Prmt5^fl/fl^* and control mice at P60. HE staining revealed no significant differences in overall brain architecture or size between groups (fig. S5A). The cerebellum of *Prmt5^fl/fl^* mice exhibited typical lobular morphology and a normal trilaminar cortex structure, including the molecular, PC, and granule cell layers (fig. S5, A to C). Nissl staining further confirmed that there were no major cytoarchitectural defects across cerebellar cortical cell types in *Prmt5^fl/fl^* mice (fig. S5D). Additionally, Calbindin immunostaining revealed normal density and spatial distribution of PCs throughout the cerebellar cortex, including both vermis and hemispheres, in *Prmt5^fl/fl^* mice at P60 (fig. S5, E to G). However, at 5 months of age, *Prmt5^fl/fl^* mice displayed extensive PC degeneration, particularly in regions associated with cerebrovascular malformations (fig. S5, H and I). These results indicate that motor deficits precede PC degeneration in *Prmt5^fl/fl^* mice.

Cerebellar motor processing is governed by the integration of excitatory and inhibitory synaptic inputs onto PCs. To explore the impact of *Prmt5* deficiency on synaptic inputs, we analyzed inhibitory and excitatory synapses targeting PCs in *Prmt5^fl/fl^*and control mice at P60. Notably, there was a significant increase in the number of positive puncta for vesicular GABA transporter (VGAT), a marker of inhibitory synapses from MLIs, in the molecular layer and surrounding PC soma of *Prmt5^fl/fl^* mice compared to controls (Fig. 2, A to C). Conversely, excitatory synapses, marked by vesicular glutamate transporter 1 (VGLUT1) in PF terminals and vesicular glutamate transporter 2 (VGLUT2) in CF terminals, exhibited comparable spatial extension or quantitative density in the molecular layer between *Prmt5^fl/fl^* and control mice (Fig. 2, D to H). This selective enhancement of MLI-PC inhibitory synaptic connectivity establishes structural evidence for functional dysregulation of PC activity, which may underlie motor deficits in *Prmt5^fl/fl^* mice.

**Figure 2.**
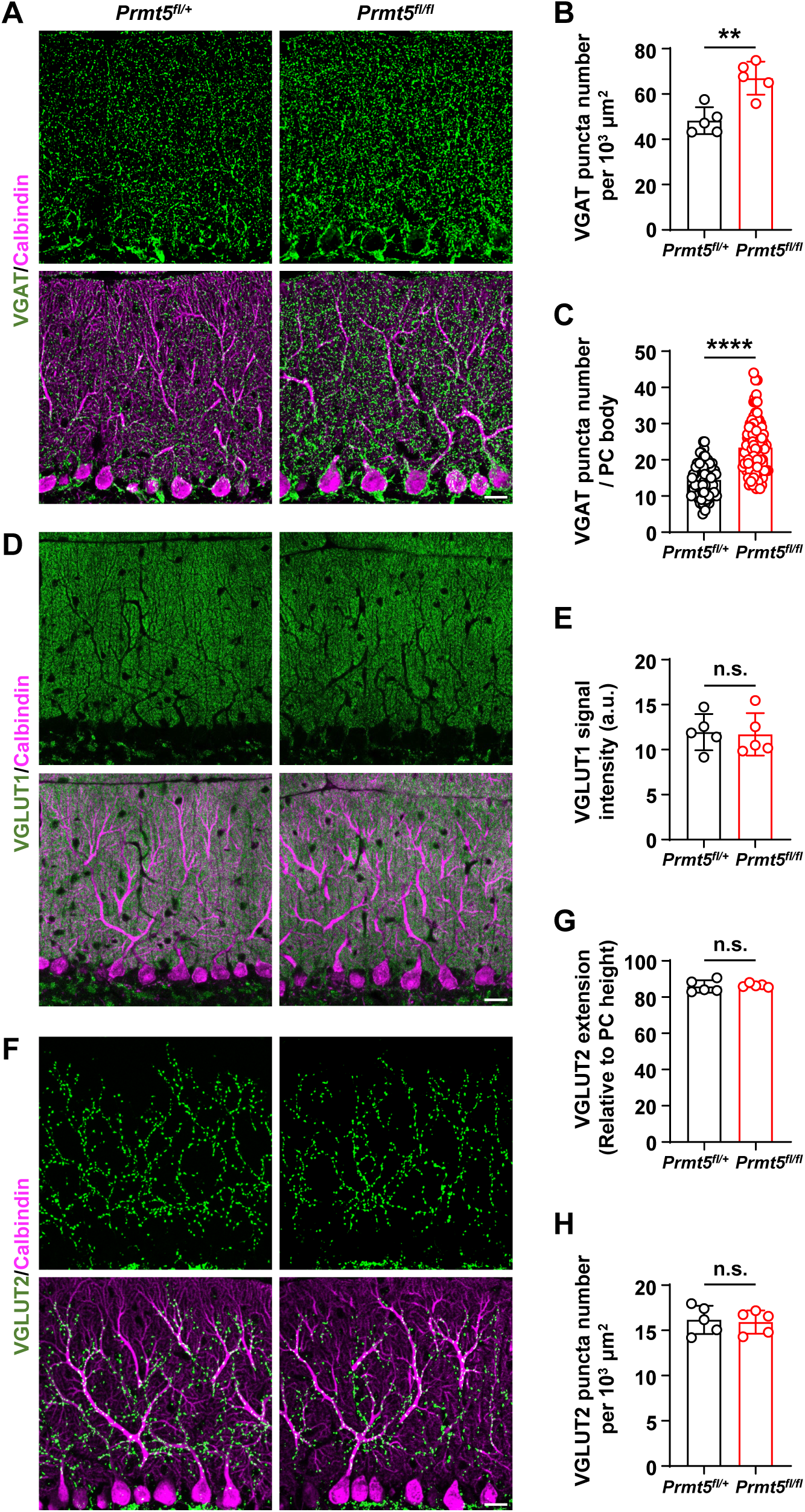
Increased inhibitory synaptic connections on PCs in *Prmt5_fl/fl_* mice. (A) Confocal images of VGAT (green) and Calbindin (purple) immunostaining in the cerebellar cortex of P60 *Prmt5_fl/+_* and *Prmt5_fl/fl_* mice. Scale bar, 20 µm. (B) Quantification of VGAT puncta number in the molecular layer. **P < 0.01 (mean ± SEM, n = 5 mice per group). (C) Quantification of VGAT puncta number surrounding individual PC bodies. ****P < 0.0001 (mean ± SEM, n = 92 PCs per group, 5 mice per group). (D) Confocal images of VGLUT1 (green) and Calbindin (purple) immunostaining. Scale bar, 20 µm. (E) Quantification of VGLUT1 signal intensity. n.s., not significant (mean ± SEM, n = 5 mice per group). (F) Confocal images of VGLUT2 (green) and Calbindin (purple) immunostaining. Scale bar, 20 µm. (G) Quantification of CF synaptic territory extension. n.s., not significant (mean ± SEM, n = 5 mice per group). (H) Quantification of VGLUT2 puncta number in the molecular layer. n.s., not significant (mean ± SEM, n = 5 mice per group).

### PC activity decreased in *Prmt5^fl/fl^* mice

To further investigate the functional consequences of increased inhibitory synapses, we recorded spontaneous inhibitory postsynaptic currents (sIPSCs) in PCs of *Prmt5^fl/fl^* and control mice at P60. *Prmt5^fl/fl^*mice exhibited a significant increase in sIPSC frequency compared to controls, while the sIPSC amplitude remained unaltered (Fig. 3, A to C). We next assessed whether augmented inhibition impairs PC activity. Cell-attached patch-clamp recordings of spontaneous action potential (sAP) firing in PCs revealed that *Prmt5^fl/fl^* exhibited a significantly lower spontaneous firing rate than controls (Fig. 3, D and E). Additionally, the coefficient of variation (CV) of the interspike intervals was significantly higher in *Prmt5^fl/fl^*PCs, indicating a disruption in the regularity of spontaneous firing activity (Fig. 3F). Collectively, these data suggest that endothelial *Prmt5* deficiency enhances inhibitory input and impairs PC activity, which may contribute to motor deficits in mice.

**Figure 3.**
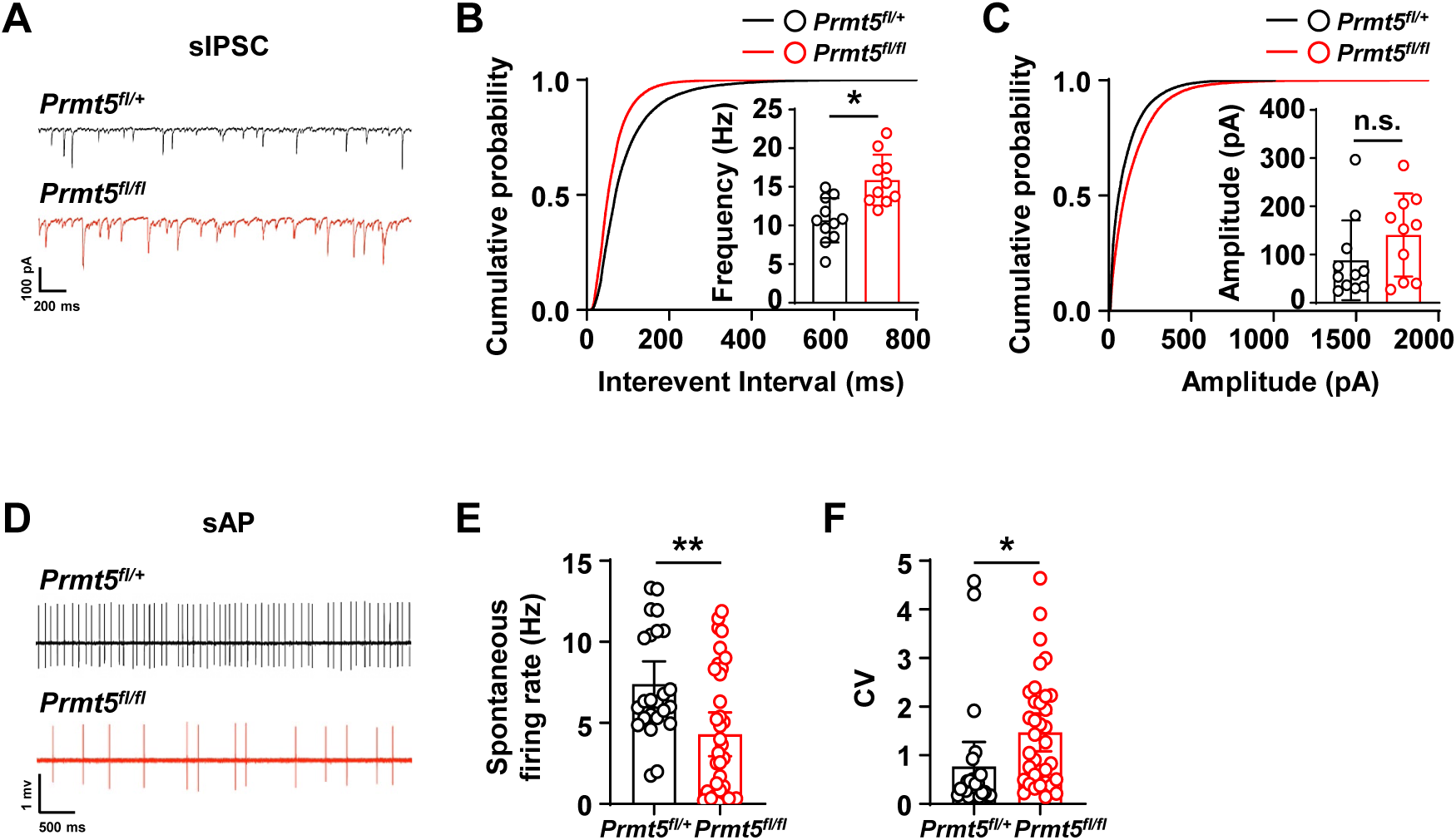
Enhanced inhibitory synaptic input and impaired intrinsic excitability in PCs of *Prmt5_fl/fl_* mice. _(A)_ Representative sIPSC traces recorded from PCs of P60 *Prmt5_fl/+_* and *Prmt5_fl/fl_* mice. (B and C) Cumulative probability distributions and quantifications of sIPSC frequency (B) and amplitude (C). *P < 0.05, n.s., not significant (mean ± SEM, n = 11 PCs in *Prmt5_fl/+_* group, n = 10 PCs in *Prmt5_fl/fl_* group, 4 mice per group). (D) Representative sAP firing traces recorded from PCs of P60 *Prmt5_fl/+_* and *Prmt5_fl/fl_* mice. (E and F) Quantifications of spontaneous firing rates (E) and coefficient of variance (CV) of interspike intervals (F). *P < 0.05, **P < 0.01 (mean ± SEM, n = 24 PCs in *Prmt5_fl/+_* group, n = 35 PCs in *Prmt5_fl/fl_* group, 4 mice per group).

### Endothelial *Prmt5* deficiency upregulates *Angptl4*

To investigate the molecular mechanisms underlying the synaptic and motor deficits in *Prmt5^fl/fl^* mice, we performed single-cell RNA sequencing (scRNA-seq) of cerebellar ECs. Fluorescence-activated cell sorting (FACS) was employed to isolate tdTomato^+^ ECs from P20 *Prmt5^fl/fl^;ROSA26^LSL-tdTomato^* and their control *Prmt5^fl/+^;ROSA26^LSL-tdTomato^* littermates, followed by droplet-based scRNA-seq utilizing the 10×Genomics platform (fig. S6A). After quality control, Seurat/SingleR analysis of 24,209 *Prmt5^fl/fl^* cells and 31,911 control cells revealed 20 distinct clusters via uniform manifold approximation and projection (UMAP) analysis (fig. S6, B and C). The proportional distribution of these clusters remained consistent between *Prmt5^fl/fl^*and controls (fig. S6D). Based on established endothelial markers, we identified 11 EC subtypes and 9 non-EC populations (fig. S6E). EC clusters were annotated as Arterial 0 (A0), Arterial 1 (A1), Capillary 0 (C0), Capillary 1 (C1), Capillary 2 (C2), Capillary 3 (C3), Capillary 4 (C4), Vein (V), Mitotic (M), Pericytes/Capillary (P/C) and Astrocytes/Capillary (Ast/C) (Fig. 4, A and B). The P/C cluster exhibited characteristics of both pericytes and capillary ECs, while the Ast/C cluster represented a mixed population of astrocytes and capillary ECs (Fig. 4, A and B). The non-ECs clusters included pericytes, fibroblast, smooth muscle cell, microglia, monocytes, ependymal cells, border-associated macrophages (BAMs), neurons, and an undefined population (Fig. 4, A and B).

**Figure 4.**
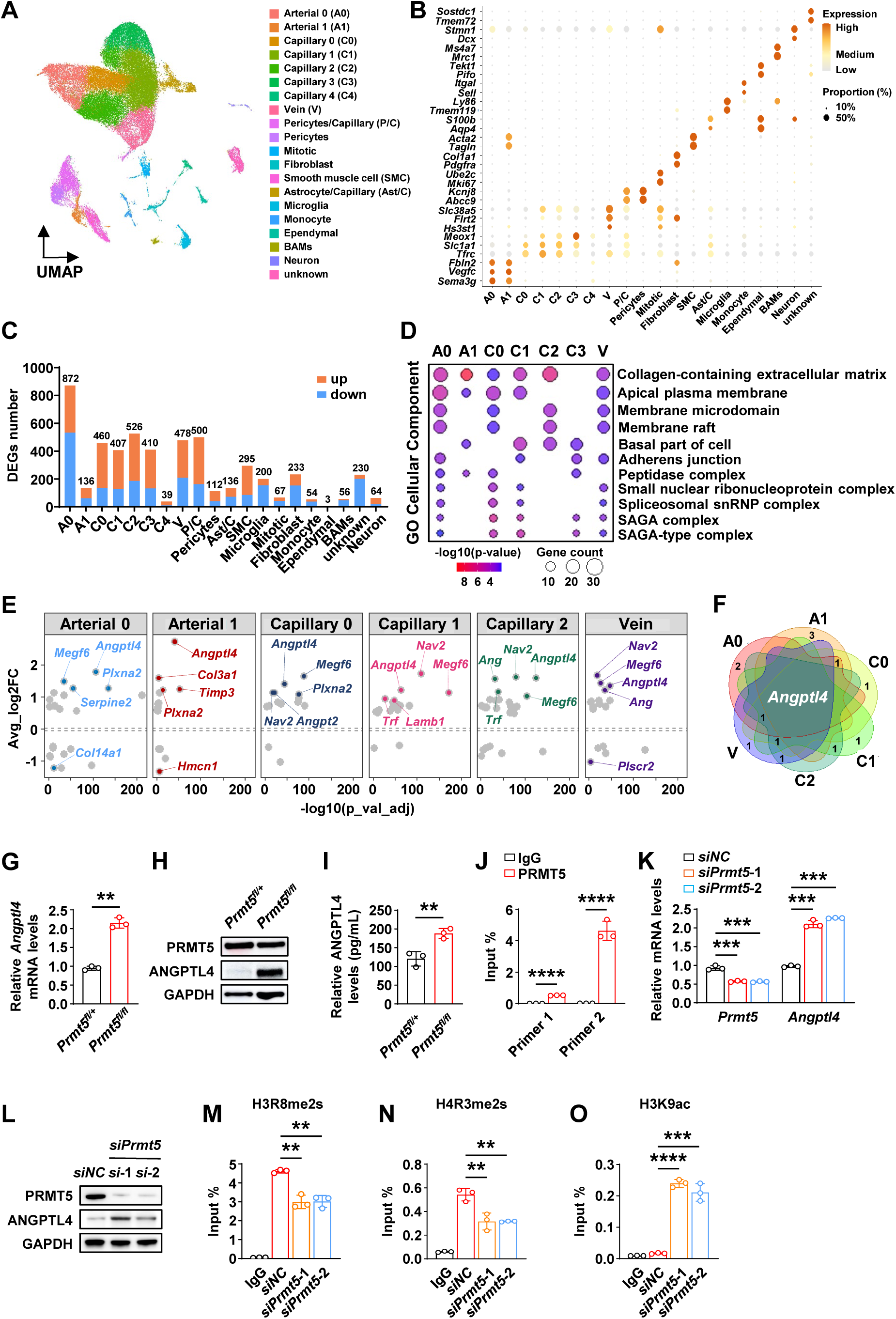
ANGPTL4 is upregulated in brain ECs of *Prmt5_fl/fl_* mice. (A) UMAP visualization of scRNA-seq data from cerebellar ECs in P20 *Prmt5_fl/+_* and *Prmt5_fl/fl_* mice, with colors corresponding to the identified clusters. (B) Dot plots showing the expression of specific marker genes in each cluster. (C) Count of significant DEGs in each cluster, with upregulation indicated in orange and downregulation in blue (average |log2FC| > 0.5, adjusted p-value < 0.05). (D) Representative gene ontology (cellular component) analysis of endothelial clusters. Dot size represents the overlap proportion between DEGs and reference gene sets; color intensity reflects -log10 (q-value). (E) Volcano plot showing DEGs encoding ECM components across 6 endothelial clusters. The top 5 genes with the greatest expression differences are labeled. (F) Venn diagram showing intersections among the top 5 genes across 6 endothelial clusters, with overlaps representing shared gene counts. _(G)_ qPCR analysis of *Angptl4* mRNA levels in brain ECs derived from P20 *Prmt5_fl/+_* and *Prmt5_fl/fl_* mice. **P < 0.01 (mean ± SEM, n = 3 per group). (H) Western blot analysis of ANGPTL4 protein expression in brain ECs derived from P20 *Prmt5_fl/+_* and *Prmt5_fl/fl_* mice. (I) ELISA analysis of secreted ANGPTL4 protein levels. **P < 0.01 (mean ± SEM, n = 3 per group). (J) ChIP-qPCR analysis of PRMT5 occupancy on the *Angptl4* promoter region in bEnd.3 cells. ****P < 0.0001 (mean ± SEM, n = 3 per group). (K) qPCR analysis of *Prmt5* and *Angptl4* mRNA levels in bEnd.3 cells transfected with *siNC* or *siPrmt5*. ***P < 0.001 (mean ± SEM, n = 3 per group). (L) Western blot analysis of PRMT5 and ANGPTL4 protein levels in bEnd.3 cells transfected with *siNC* or *siPrmt5*. GAPDH served as the loading control. (M-O) ChIP-qPCR analysis of the enrichment of H3R8me2s (M), H4R3me2s (N) and H3K9ac (O) at the *Angptl4* promoter region in bEnd.3 cells transfected with *siNC* or *siPrmt5*. **P < 0.01, ****P < 0.0001 (mean ± SEM, n = 3 per group).

To determine the effect of *Prmt5* deficiency, we examined differentially expressed genes (DEGs) across EC clusters, applying a threshold of |log2FC| > 0.5 and an adjusted p-value < 0.05. Significant transcriptomic alterations were observed in A0, C0, C1, C2, C3 and V clusters, whereas the C4 population remained largely unaffected (Fig. 4C). Notably, GO enrichment analysis identified significant alterations in endothelial-secreted collagen-containing extracellular matrix (ECM) components across A0, A1, C0, C1, C2, and V subpopulations, with at least 14 DEGs implicated (Fig. 4D). The ECM constitutes a dynamic, three-dimensional macromolecular network comprising diverse biomolecules, including an estimated 300 distinct proteins such as collagens, glycoproteins, and proteoglycans, along with both secreted and cell-bound components (*41, 42*). Beyond providing structural support, the ECM plays an essential role in regulating cellular behavior by mediating bidirectional (inside-out and outside-in) signaling, which orchestrates complex signaling pathways that govern cellular function, phenotype, morphology, and behavior (*43, 44*). We therefore hypothesized that PRMT5-mediated alterations in endothelial secreted ECM components might contribute to excessive inhibitory synaptic connections in the cerebellum. By profiling genes associated with collagen-containing ECM components, we identified significant *Angptl4* upregulation across these endothelial clusters (Fig. 4, E and F). ANGPTL4, a multifunctional secreted protein, has pivotal impacts on ECs in lipid metabolism (*45*), angiogenesis (*46*), and vascular permeability (*47*). Previous studies have shown that ANGPTL4 expression is elevated in various cerebrovascular diseases, such as ischemic stroke (*48*) and atherosclerosis (*49*). However, its physiological roles in the synapse formation and motor coordination remain largely unclear. Our data suggest that ANGPTL4 may serve as a pivotal mediator in PRMT5-regulated endothelial dysfunction. Consistently, both mRNA and protein levels of *Angptl4* were significantly upregulated in brain ECs of *Prmt5^fl/fl^* mice (Fig. 4, G and H), and ELISA confirmed elevated ANGPTL4 secretion in culture supernatants of primary brain ECs from *Prmt5^fl/fl^*mice (Fig. 4I).

To elucidate the mechanism by which PRMT5 regulates *Angptl4* transcription, we examined its role in epigenetic regulation. We conducted chromatin immunoprecipitation (ChIP) assays in brain microvascular endothelial bEnd.3 cells to determine whether PRMT5 directly modulates *Angptl4* transcription. ChIP-qPCR results demonstrated that PRMT5 binds directly to the *Angptl4* promoter (Fig. 4J). Based on the established role of PRMT5 in mediating symmetric dimethylation of histones, we hypothesized that it represses *Angptl4* expression through H3R8me2s and H4R3me2s modifications, which are recognized as repressive marks for gene expression (*50, 51*). To test this, we performed RNA interference in bEnd.3 cells and observed an approximately two-fold upregulation of *Angptl4* transcription in *siPrmt5*-treated cells compared to *siNC* controls (Fig. 4, K and L). Subsequent ChIP-qPCR assays revealed that PRMT5 knockdown reduced the enrichment of H3R8me2s and H4R3me2s at the *Angptl4* promoter (Fig. 4, M and N). Concomitantly, a marked increase in H3K9 acetylation (H3K9ac) at the promoter was detected upon PRMT5 knockdown (Fig. 4O), indicating a more transcriptionally permissive chromatin state (*52*). Consistent with these findings, similar alterations in histone modifications were recapitulated in NIH-3T3 cells (fig. S7). These results provide compelling evidence that PRMT5 represses *Angptl4* transcription through symmetric dimethylation of H3R8 and H4R3.

Together, these data demonstrate that *Prmt5* deficiency in cerebrovascular ECs leads to the derepression of *Angptl4*, which may contribute to increased inhibitory synapses in *Prmt5^fl/fl^* mice.

### ANGPTL4 promotes inhibitory synaptogenesis

We further investigated the effect of elevated ANGPTL4 on inhibitory synaptogenesis. We first co-cultured the *siPrmt5* or *siNC* bEnd.3 cells with the primary neurons isolated from embryonic day 13.5 (E13.5) mouse embryos. Quantitative immunofluorescence revealed a significant increase in VGAT^+^ puncta density along neuronal dendrites in *siPrmt5* co-cultures compared to *siNC* controls (Fig. 5, A and B), indicating that *Prmt5*-deficient ECs promotes inhibitory synaptogenesis via secreted factors.

**Figure 5.**
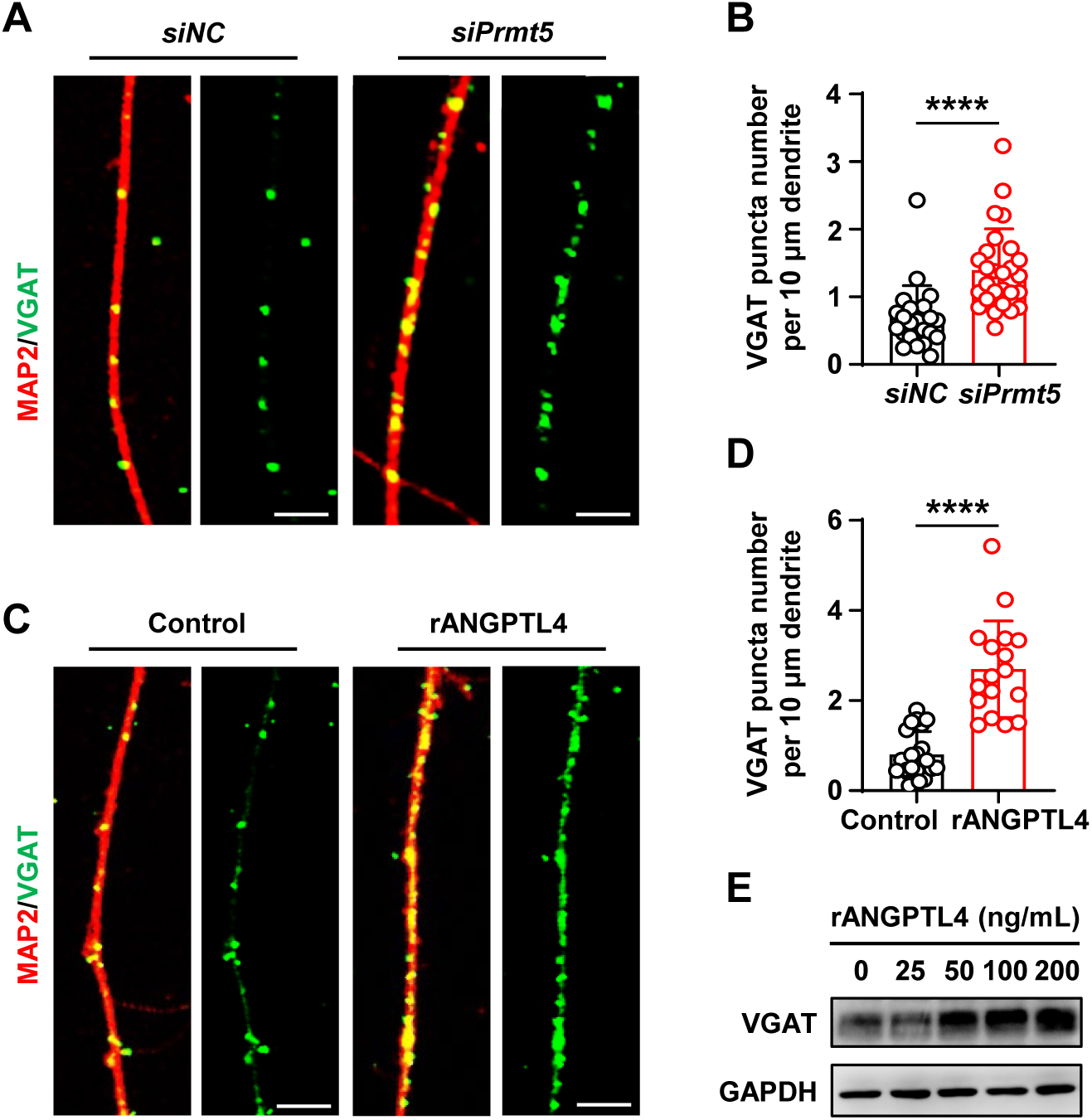
ANGPTL4 promotes inhibitory synapse formation. (A) Confocal images of VGAT (green) and MAP2 (red) immunostaining in neurons co-cultured with bEnd.3 cells transfected with *siNC* or *siPrmt5*. Scale bar, 5 µm. (B) Quantification of VGAT_+_ inhibitory synapses in (A). ****P < 0.0001 (mean ± SEM, n = 22 cells in *siNC* group, n = 27 cells in *siPrmt5* group, 3 independent experiments). (C) Confocal images of VGAT (green) and MAP2 (red) immunostaining in primary neurons dissociated from E13.5 mouse embryos treated with 0 ng/mL (control) or 50 ng/mL rANGPTL4. Scale bar, 5 μm. (D) Quantification of VGAT_+_ inhibitory synapses in (C). ****P < 0.0001 (mean ± SEM, n = 22 cells in control group, n = 17 cells in rANGPTL4 group, 3 independent experiments). (E) Western blot analysis of VGAT expression in neurons treated with the indicated doses of rANGPTL4. GAPDH served as the loading control.

We next treated the primary neurons dissociated from E13.5 mouse embryos with recombinant ANGPTL4 protein (rANGPTL4). Immunofluorescence analysis demonstrated a significant elevation in VGAT^+^ puncta density on neuronal cells after rANGPTL4 administration (Fig. 5, C and D). Additionally, western blot analysis revealed a dose-dependent upregulation of VGAT protein expression corresponding to increasing rANGPTL4 concentrations (Fig. 5E). These results collectively demonstrate that elevated EC-derived ANGPTL4 facilitates inhibitory synaptogenesis.

### Deletion of *Angptl4* in cerebrovascular ECs normalizes inhibitory synaptic input onto PCs and rescues motor deficits in *Prmt5^fl/fl^*mice

To determine whether elevated ANGPTL4 is responsible for the increased inhibitory synapses and motor deficits in *Prmt5^fl/fl^* mice, we generated *Prmt5* and *Angptl4* double-knockout mice. The *Angptl4^flox/flox^*mice were generated by deleting exons 4 to 6 in the *Angptl4* locus (fig. S8A). *Angptl4^flox/flox^* mice were then bred with *SP-A-Cre*;*Prmt5^fl/+^* mice to generate *SP-A-Cre*;*Prmt5^fl/+^;Angptl4^fl/+^*(*Prmt5^fl/+^;Angptl4^fl/+^*), *SP-A-Cre*;*Prmt5^fl/+^;Angptl4^fl/fl^* (*Prmt5^fl/+^;Angptl4^fl/fl^*), *SP-A-Cre*;*Prmt5^fl/fl^;Angptl4^fl/+^* (*Prmt5^fl/fl^;Angptl4^fl/+^*), and *SP-A-Cre*;*Prmt5^fl/fl^;Angptl4^fl/fl^* (*Prmt5^fl/fl^;Angptl4^fl/fl^*) mice. qPCR showed effective deletion of *Angptl4* gene in brain ECs of *Prmt5^fl/+^;Angptl4^fl/fl^*mice compared to controls (fig. S8B).

We next examined the inhibitory synapse density and electrophysiological characteristics of PCs in *Prmt5^fl/fl^;Angptl4^fl/fl^*mice at P60. Notably, the density of VGAT^+^ inhibitory synapses in both the molecular layer and around PC soma in *Prmt5^fl/fl^;Angptl4^fl/fl^*mice significantly deceased compared to *Prmt5^fl/fl^;Angptl4^fl/+^*mice (Fig. 6, A to C). Consistently, electrophysiological recordings showed that the frequency of sIPSC in *Prmt5^fl/fl^;Angptl4^fl/fl^* mice was restored to levels comparable to those in *Prmt5^fl/+^;Angptl4^fl/+^*controls (Fig. 6, D to F).

**Figure 6.**
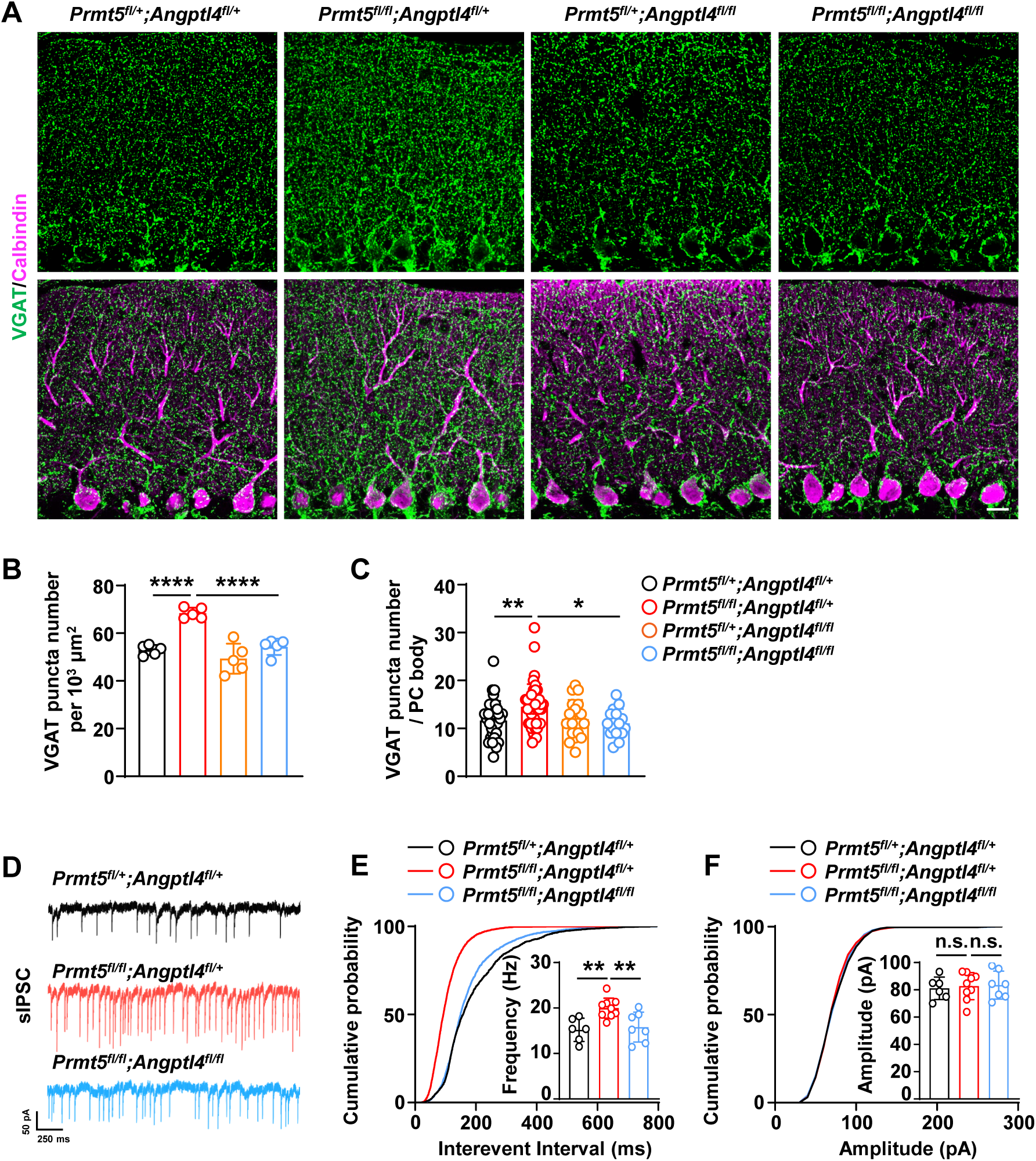
Cerebrovascular *Angptl4* knockout rescues abnormal inhibitory synaptic input onto PCs in *Prmt5_fl/fl_* mice. (A) Confocal images of VGAT (green) and Calbindin (purple) immunostaining in the cerebellar cortex of P60 *Prmt5_fl/+_;Angptl4_fl/+_*, *Prmt5_fl/fl_;Angptl4_fl/+_*, *Prmt5_fl/+_;Angptl4_fl/fl_*, and *Prmt5_fl/fl_;Angptl4_fl/fl_* mice. Scale bar, 20 µm. (B and C) Quantification of VGAT puncta number in the molecular layer (B) and surrounding individual PC bodies (C). *P < 0.05, **P < 0.01, ****P < 0.0001 (mean ± SEM, n = 5 mice per group). (D) Representative sIPSC traces recorded from PCs of P60 *Prmt5_fl/+_;Angptl4_fl/+_*, *Prmt5_fl/fl_;Angptl4_fl/+_*, and *Prmt5_fl/fl_;Angptl4_fl/fl_* mice. (E and F) Cumulative probability distributions and quantifications of sIPSC frequency (E) and amplitude (F). **P < 0.01, n.s., not significant (mean ± SEM, n = 6 PCs in *Prmt5_fl/+_;Angptl4_fl/+_* group, n = 9 PCs in *Prmt5_fl/fl_;Angptl4_fl/+_* group, n = 7 PCs in *Prmt5_fl/fl_;Angptl4_fl/fl_* group, 4 mice per group).

Finally, we assessed whether knockout *Angptl4* could ameliorate motor deficits in *Prmt5^fl/fl^* mice. Remarkably, *Prmt5^fl/fl^;Angptl4^fl/fl^*mice exhibited significant improvements in motor performance (Fig. 7). In the footprint analysis, stride lengths were notably increased in *Prmt5^fl/fl^;Angptl4^fl/fl^*mice compared to *Prmt5^fl/fl^;Angptl4^fl/+^* littermates, although they remained slightly shorter than those of *Prmt5^fl/+^;Angptl4^fl/+^* controls (Fig. 7A). Similarly, in the rotarod and balance beam tests, *Prmt5^fl/fl^;Angptl4^fl/fl^*mice exhibited enhanced motor coordination, as indicated by prolonged rotarod latency and shorter balance beam traversal time relative to *Prmt5^fl/fl^;Angptl4^fl/+^*mice (Fig. 7, B and C).

**Figure 7.**
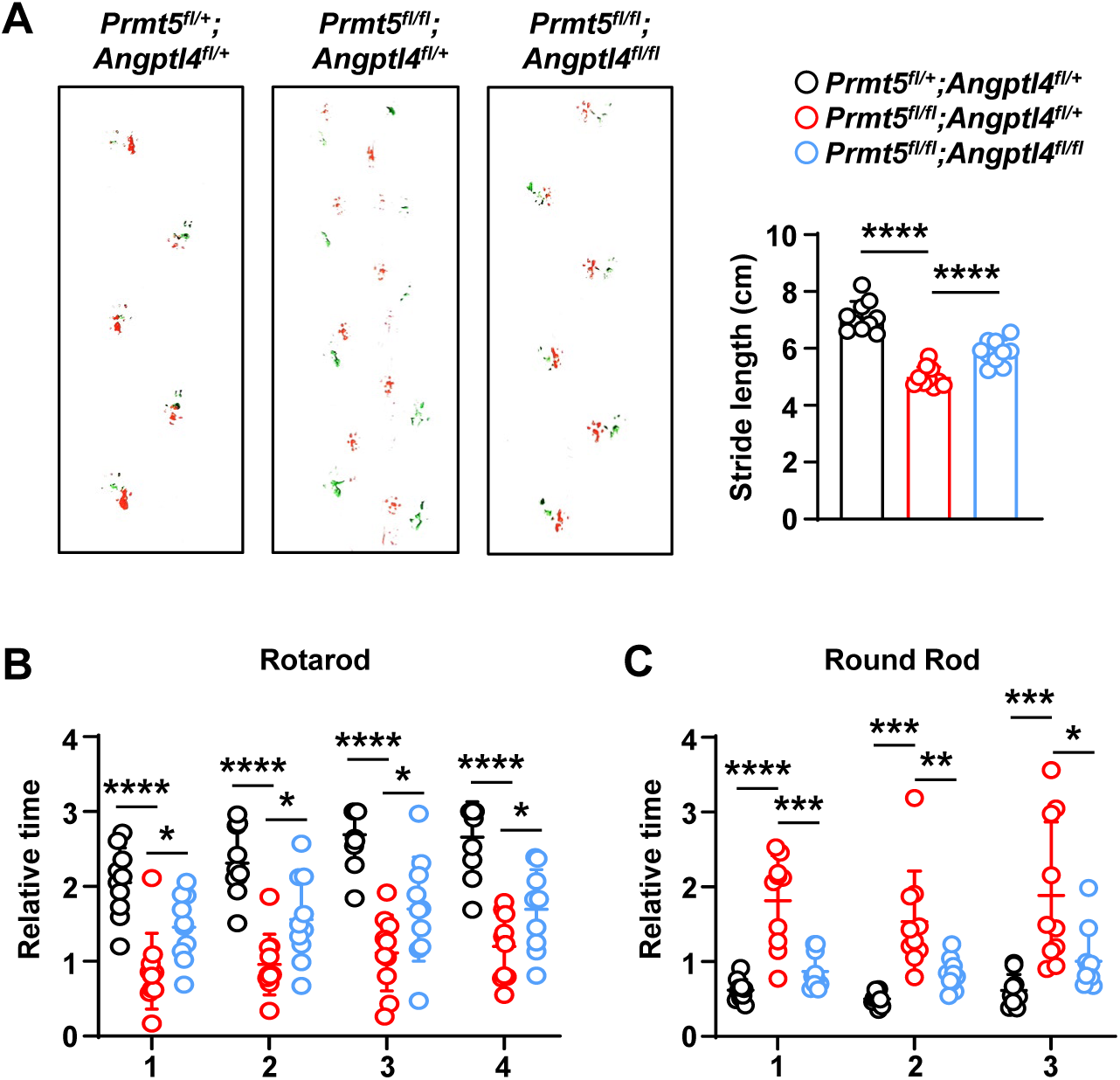
Cerebrovascular *Angptl4* knockout relieves the motor deficits in *Prmt5_fl/fl_* mice. (A) Representative footprints and quantification of stride length in P60 *Prmt5_fl/+_;Angptl4_fl/+_*, *Prmt5_fl/fl_;Angptl4_fl/+_*, and *Prmt5_fl/fl_;Angptl4_fl/fl_* mice. Red, forepaws; green, hindpaws. ****P < 0.0001 (mean ± SEM, n = 10 mice per group). (B) Quantification of latency time spent on the rotarod. Data are from four independent trials. *P < 0.05, ****P < 0.0001 (mean ± SEM, n = 10 mice per group). (C) Quantification of time spent crossing the balance beam. Data are from three independent trials. *P < 0.05, **P < 0.01, ***P < 0.001, ****P < 0.0001 (mean ± SEM, n = 10 mice per group).

Collectively, these results demonstrate that *Angptl4* deletion effectively rescues the synaptic abnormalities and motor deficits in *Prmt5^fl/fl^*mice, supporting the notion that ANGPTL4 upregulation mediates, at least in part, the motor dysfunction induced by PRMT5 deficiency .

## Discussion

The role of the cerebrovasculature in the development and modulation of neural circuitry remains largely unexplored. In this study, we uncover a novel mechanism by which brain ECs control inhibitory synaptogenesis and motor coordination through the PRMT5-ANGPTL4 axis, demonstrating that cerebrovascular dysfunction can directly disrupt motor function through synaptic dysregulation.

Our findings reveal a previously unrecognized physiological role of brain ECs in modulating inhibitory synaptogenesis in the cerebellum. While brain ECs are increasingly recognized for their roles in neurodevelopment and synaptic plasticity (*5, 7*), their function in cerebellar synaptic organization has remained enigmatic. Here, we demonstrate that endothelial PRMT5 critically maintains cerebellar circuit homeostasis by repressing ANGPTL4, an angiocrine factor that drives excessive inhibitory synaptogenesis onto PCs. In mice lacking endothelial PRMT5, ANGPTL4 upregulation led to increased inhibitory input onto PCs, suppressed PC firing, and severe motor deficits, indicating that the endothelial PRMT5-ANGPTL4 axis preserves the excitatory-inhibitory balance essential for motor coordination. Supportively, cerebrovascular-specific deletion of *Angptl4* normalizes inhibitory synaptic input onto PCs and rescues motor deficits in *Prmt5*-deficient mice. This finding challenges the prevailing view that cerebrovascular contributions to motor impairment are primarily mediated by hemodynamic or BBB dysfunction (*53, 54*). Instead, our data support a model in which brain ECs actively shape neural circuits through paracrine mechanisms (*4–7*), with PRMT5-ANGPTL4 axis serving as a key regulator of this process.

A key discovery of this study is the identification of ANGPTL4 as an endothelial-derived synaptogenic angiocrine factor. While ANGPTL4 has been implicated in neuroprotection after ischemic stroke (*55, 56*) and in Parkinson’s models (*57*), it has also been linked to pathological processes such as BBB disruption, neuronal apoptosis, and microglial dysfunction (*58–61*). Despite these diverse roles, a function in synapse regulation had not been reported. Here, we demonstrate that endothelial-derived ANGPTL4 promotes inhibitory synaptogenesis and that its dysregulation leads to circuit dysfunction. Mechanistically, endothelial *Prmt5* deficiency promotes *Angptl4* transcription by reducing symmetric dimethylation of H3R8 and H4R3 and enhancing H3K9 acetylation at its promoter. These findings establish a direct link between endothelial epigenetic regulation and cerebellar circuit function.

Our findings redefine the mechanistic link between cerebrovascular dysfunction and motor impairment, offering new insights into the pathogenesis of vascular-related motor disorders. Although clinical studies have linked cerebrovascular disorders to gait and balance disturbances (*62, 63*), whether vascular cells actively regulate motor coordination at the synaptic level has remained unknown. Here, we demonstrate that endothelial defects can drive motor deficits via synaptic dysregulation, independent of overt vascular damage. Notably, motor impairments emerged as early as P20 in our model, preceding any detectable alterations in cerebellar vasculature. This temporal dissociation indicates that synaptic dysfunction initiated by endothelial dysfunction is sufficient to cause motor impairment. This finding opens new avenues for therapeutic intervention, as targeting endothelial-derived factors such as ANGPTL4 could restore synaptic balance without requiring vascular repair. Indeed, brain endothelial-specific ablation of *Angptl4* significantly rescued motor deficits in *Prmt5*-deficient mice, demonstrating the feasibility of vascular-targeted therapies for synaptic disorders. That the rescue was incomplete suggests additional endothelial-derived factors contribute to the phenotype, a possibility supported by our scRNA-seq data and deserving of further investigation.

In summary, this study identifies a previously unrecognized endothelial-to-neural signaling axis essential for motor coordination, providing a new conceptual framework for vascular-focused therapeutic strategies, not only for motor disorders but also for other neurodegenerative conditions associated with cerebrovascular dysregulation.

## Materials and Methods

### Experimental Design

Animals were randomly assigned to experimental groups, with experimenters blinded to genotype information throughout the study duration. No statistical methods were used to predetermine the sample size. In conformity to the 3Rs (Reduction, Replacement and Refinement), the smallest sample size that would give a significant difference was used. No data or animals were excluded from the analysis.

### Mice

All mice were housed and bred in a specific-pathogen-free (SPF) facility accredited by the Laboratory Animal Management Committee (LAMC) of the Chinese People’s Liberation Army (CPLA). Mice were maintained under a 12-hour reverse light/dark cycle (lights on at 8:00 am) with controlled temperature and humidity. All animal procedures were performed in strict compliance with the ethical guidelines and protocols approved by the Institutional Animal Care and Use Committee (IACUC) at the Beijing Institute of Lifeomics.

*Prmt5^flox/flox^* (*Prmt5^fl/fl^*) mice and *SP-A-Cre* transgenic mice were previously described. *Angptl4^flox/flox^* (*Angptl4^fl/fl^*) mice were purchased from Cyagen Biosciences. To selectively knockout *Prmt5* in endothelial cells, *SP-A-Cre* transgenic mice were crossed to *Prmt5^fl/fl^*mice to generate *SP-A-Cre*;*Prmt5^fl/fl^* mice. To trace cerebral endothelial cells and monitor Cre-mediated recombination, *SP-A-Cre*;*Prmt5^fl/fl^* mice were bred with *ROSA26^LSL-tdTomato^* transgenic mice. *SP-A-Cre*;*Prmt5^fl/fl^* mice were further crossed to *Angptl4^fl/fl^* mice to obtain

*Prmt5* and *Angptl4* double knockout mice (*SP-A-Cre*;*Prmt5^fl/fl^*;*Angptl4^fl/fl^*). In this study, heterozygous mice served as the littermate control and did not exhibit the motor deficit phenotype. Only male mice were included and randomly allocated to experimental groups. The ages of mice used in experiments are specified in the respective sections of the text.

### Cell culture

Mouse embryonic fibroblasts (NIH-3T3 cells, ATCC, CRL-1658) and mouse brain endothelial cells (bEnd.3 cells, ATCC, CRL-2299) were cultured at 37 °C in a humidified atmosphere containing 5% CO₂. All cell lines were obtained directly from the American Type Culture Collection (ATCC) and used without further authentication. Cells were maintained in DMEM (VivaCell, C3113-0500) supplemented with 10% fetal bovine serum (FBS) (VivaCell, C04001-500) and 1% penicillin-streptomycin (VivaCell, C3420-0100).

### Multiplex immunofluorescence assay

Mice were deeply anesthetized with tribromoethanol and transcardially perfused with saline followed by 4% paraformaldehyde (PFA). Brains were post-fixed in 4% PFA at 4 °C overnight, then either dehydrated through graded ethanol series and embedding in paraffin, or equilibrated in 30% sucrose and embedded in NEG-50 medium for cryosectioning. Paraffin-embedded tissues were sectioned at 6 μm and stored at room temperature, while frozen tissues were sectioned at 45 μm and preserved in cryoprotectant solution (glycerol, ethylene glycol, and 0.1 M phosphate buffer, pH 7.4, 1:1:2 by volume) at -20 °C.

For antigen retrieval, paraffin sections were microwaved in citrate buffer (pH 6.0) at boiling temperature for 20 seconds and cooled at room temperature. Frozen sections were permeabilized with TBS++ (TBS containing 3% donkey serum and 0.3% Triton X-100) for 4 hours at room temperature. Endogenous peroxidase activity was quenched with 3% hydrogen peroxide for 15 minutes, followed by blocking with goat serum (ZSGB-BIO, ZLI-9022) for 30 minutes at 37 °C. Primary antibodies (listed below) were incubated overnight at 4 °C. HRP-conjugated secondary antibodies were applied for 30 minutes at 37 °C, followed by Tyramide Signal Amplification (TSA)-fluorophores (Histova Biotechnology, NECC7100) for fluorescence detection. For multiple labeling, sections were reheated in retrieval/elution buffer (Histova Biotechnology, ABCFR5L), and the staining procedure was repeated with different fluorophores. The antibodies were listed in Supplementary Table 1.

### Immunocytochemistry

Primary cortical neurons cultured on poly-L-lysine-coated coverslips in 24-well plates were fixed with 4% PFA (15 minutes) and permeabilized with 0.5% Triton X-100. After blocking with goat serum, cells were incubated overnight at 4 °C with anti-MAP2 (GeneTex, GTX634473) and anti-VGAT (GeneTex, GTX101908). Fluorescence-conjugated secondary antibodies (Proteintech, RGAR004 and RGAM005) were applied for 1 hour at 37 °C. Coverslips were mounted with antifade medium for confocal imaging. The antibodies were listed in Supplementary Table 1.

### Confocal microscopy and image processing

Confocal imaging was performed using a Zeiss LSM 880 laser scanning confocal microscope at room temperature. Images were acquired using the following objectives: EC Plan-Neofluar 10x/0.30 M27, EC Plan-Neofluar 20x/0.50 M27, EC Plan-Neofluar 40x/0.75 M27, and Plan-Apochromat 63x/1.4 Oil DIC M27. For co-localization analysis of cell markers, 3-dimensional Z-stack images were acquired. Z-stack alignments were processed using ZEN software (Carl Zeiss). To image the entire cerebellum, tile scans were performed at 20x magnification with 10% spatial overlap between adjacent tiles. Tiles were automatically stitched and assembled using ZEN software (Carl Zeiss).

### Nissl and HE staining

For Nissl staining, sections from the cerebellar vermis were stained with Nissl Staining solution (Beyotime, C0117) for 5 minutes at room temperature according to the manufacturer’s instructions. For HE staining, sections from the cerebellar vermis were stained with hematoxylin (ZSGB-BIO, ZLI-9610) for 90 seconds at room temperature. After differentiation and bluing, sections were counterstained with eosin (ZSGB-BIO, ZLI-9613) for 20 seconds at room temperature. Stained sections were imaged using a light microscope (Carl Zeiss, Axio Imager 2) under bright-field illumination.

### Western blot analysis

Cellular proteins were extracted using RIPA buffer (50 mM Tris-HCl pH 8.0, 150 mM NaCl, 5 mM EDTA, 0.1% SDS, and 1% NP-40) supplemented with protease inhibitors (Roche, 4693159001) and phosphatase inhibitors (Roche, 4906837001). Lysates were centrifuged at 16,000 × g for 10 minutes at 4 °C, and the supernatant was collected. Protein concentrations were quantified using the BCA Protein Assay Reagent (Thermo Fisher Scientific, 23225). Equivalent quantities of protein samples (20 μg protein) were separated by SDS-PAGE and transferred onto PVDF membranes (Immobilon-P, IPVH00010). Membranes were blocked with 10% skim milk for 1 hour at room temperature and then incubated with primary antibodies overnight at 4 °C. After washing, membranes were incubated with peroxidase-conjugated secondary antibodies: Goat anti-Rabbit IgG (ZSGB-BIO, ZB-2301) or Goat anti-Mouse IgG (ZSGB-BIO, ZB-2305). Signals were detected using the Enlight detection system (Engreen Biosystem, 29100), and densitometry was performed with ImageJ, using GAPDH as a loading control. The antibodies were listed in Supplementary Table 1.

### qPCR

Total RNA was extracted using Trizol reagent (Invitrogen, 15596026) and reverse transcribed into cDNA using ReverTra Ace® qPCR RT Kit (TOYOBO, FSQ-201). qPCR was performed using SYBR Green PCR Master Mix (TOYOBO, QPK-201) on a QuantStudio 3 Real-Time PCR system (Thermo Fisher). Gene expression levels were normalized to the housekeeping gene *Gapdh*. Primer sequences used in this study were as following: *Gapdh*, 5’-CATCACTGCCACCCAGAAGACTG-3’ and 5’-ATGCCAGTGAGCTTCCCGTTCAG-3’; *Prmt5*, 5’-CCTGCTTTACCTTCAGCCATCC-3’ and 5’-GCACAGTCTCAAAGTAGCCTGC-3’; *Angptl4*, 5’-CTGGACAGTGATTCAGAGACGC-3’ and 5’-GATGCTGTGCATCTTTTCCAGGC-3’.

### Chromatin immunoprecipitation (ChIP) assay

Briefly, 10^7^ cells were cross-linked with 1% formaldehyde for 10 minutes at room temperature, and the reaction was quenched by adding 0.125 M glycine for 5 minutes. Cells were then washed in PBS, resuspended in ChIP lysis buffer (10 mM Tris pH 8.0, 150 mM NaCl, 0.1% SDS, 1 mM EDTA, and protease inhibitors), and sonicated using a VCX750 sonicator (Sonics) to shear chromatin into fragments of 200–500 bp. Immunoprecipitation was performed using either control IgG or specific antibodies targeting the protein of interest. Precipitated DNA was purified and analyzed by qPCR. The antibodies were listed in Supplementary Table 1. Primer sequences targeting the *Angptl4* promoter were as follows: Primer 1 5’-ACGCAGGACACAGGGGAATA-3’ and 5’-GACCAGGCTGACCTCAAACT-3’ Primer 2 5’-GTAGGGCAGAGTTACGTGGC-3’ and 5’-GGTGGTGGAAGTGAGAGTGG-3’.

### Cell isolation and culture

Brain ECs used in this study were isolated from *Prmt5^fl/+^* and *Prmt5^fl/fl^* mice. Brains were minced and enzymatically digested in a solution containing 1 mg/mL collagenase I (Sigma, C0130) and DNase I (Roche, 10104159001) at 37 °C for 30 minutes, with intermittent pipetting to enhance tissue dissociation. The digested tissue was centrifuged at 350 × g for 10 minutes, and the cell pellet was resuspended in 20% bovine serum albumin (BSA) solution. After centrifugation and myelin removal, the dissociated cells were incubated with biotin-conjugated anti-mouse CD31 antibody (BioLegend,102503) for 30 minutes at 4 °C to label ECs. Cells were washed with 2% FBS/PBS and further incubated with streptavidin-conjugated magnetic particles (BD, 557812) for 30 minutes at 4 °C. The labeled cells were centrifuged, resuspended in 2% FBS/PBS, and passed through MS columns (Miltenyi Biotec, 130-042-901) to enrich CD31^+^ cells. The CD31^+^ cell fraction was collected by centrifugation at 2000 × g for 5 minutes at 4 °C. Isolated primary ECs were utilized for culture or for the detection of RNA or protein expression.

Cortical neurons were dissociated from E13.5 mice embryos. Brain tissues were dissociated using StemPro^TM^ Accutase^TM^ cell dissociation reagent (GIBCO, A1110501) for 4 minutes at 37 °C. Dissociated neurons were plated on round coverslips (Electron Microscopy Science, 72196-12) pre-coated with poly-L-lysine (Pricella, PB180523) and cultured in DMEM supplemented with 20% (v/v) FBS. After 6 hours of plating, the medium was replaced with neurobasal medium (GIBCO, 21103049) supplemented with B27 (GIBCO, 17504-044) and GlutaMAX (GIBCO, 35050-061), with the change of medium every third day.

For the bEnd.3-neuron co-culture system, primary neurons (150,000 cells/well) were initially plated on poly-L-lysine-coated glass coverslips in 24-well plates and cultured for 72 hours. Transwell inserts (Corning Falcon, 353104) pre-seeded with 80,000 bEnd.3 cells transfected with *siNC* or *siPrmt5* were then transferred into the neuron-containing plates. After 96 hours of co-culture, neurons were fixed with 4% PFA for subsequent immunostaining, confocal imaging, and quantitative analysis.

## ANGPTL4 ELISA

Isolated brain ECs were cultured in EGM-2 medium (Lonza Bioscience, CC-3162), with the medium replaced every three days. To measure extracellular ANGPTL4 protein levels, culture medium collected from day 4 to day 6 was analyzed using a Mouse Angiopoietin-like 4 (ANGPTL4) ELISA Kit (CUSABIO, CSB-EL001712MO) according to the manufacturer’s instructions.

### RNAi and cell transfection

Short interfering RNAs (siRNAs) targeting the mouse *Prmt5* gene were synthesized by Tsingke (Beijing). Transfection of *siPrmt5* or negative control *siNC* into cells was performed using jetPRIME (Polyplus, 101000046) according to the manufacturer’s instructions. Cells were harvested 48 hours after transfection for further analysis. The siRNA sequences used were as followings: *siPrmt5*-1, sense-GCACAGUUUGAGAUGCCUU; *siPrmt5*-2, sense-GACAACAACCGCUACUGUA; *siNC*, sense-UUCUCCGAACGUGUCACGUTT.

### Behavioral tests

All behavioral assessments were conducted on group-housed male mice by experimenters blinded to genotype.

### Footprint assay

Forepaws and hindpaws were marked with non-toxic red and green ink, respectively. Mice traversed a narrow tunnel lined with white paper, and stride length was measured as the distance between consecutive ipsilateral paw prints. Only the clear and sequential footprints with ≥ 2 uninterrupted steps were analyzed.

### Balance beam test

A 75-cm-long beam (11 mm width) was elevated 50 cm above ground at a 30° incline, with a dark shelter positioned at the elevated end. Mice underwent two training trials (navigating from the lower to the higher end) followed by three consecutive test trials. The time to traverse the full beam length was recorded.

### Rotarod test

Before each trial, mice were trained on a rotating drum (3 cm in diameter, Ugo Basile, 47650) at a slow speed (4 rpm over 3 minutes). The rod was then accelerated from 4 to 40 rpm over 5 minutes, and the latency to fall was recorded. If a mouse clung to the rod and completed three full rotations, it was scored as fallen at the time. Each mouse was tested four times daily for 4 consecutive days, with a 20 to 30 minutes interval between trails.

### Ex vivo electrophysiology

For ex vivo analysis of isolated but functional neuronal networks, cerebellums from P60 mice were used. Animals were anesthetized by inhalation of 5% (v/v) isoflurane and rapidly decapitated. The whole cerebellum was then removed and sliced (300 μm). Brain slices were incubated in oxygenated artificial cerebrospinal fluid (aCSF) containing 5 mM KCl, 124 mM NaCl, 1.25 mM NaH_2_PO_4_, 10 mM glucose, 26 mM NaHCO_3_, 2.4 mM CaCl_2_, and 1.2 mM MgCl_2_ (pH 7.3–7.5, osmolarity 292–320 mOsm), continuously bubbled with a gas mixture of 95% O_2_ and 5% CO_2_. All recordings were obtained from the soma of PCs in cerebellar vermis slices. PCs were visualized using a standard upright microscope (BX51WI, Olympus).

For spontaneous inhibitory postsynaptic current (sIPSC) recordings, patch electrodes were filled with an internal solution containing 140 mM KCl, 10 mM HEPES, 4 mM MgCl_2_, 0.1 mM EGTA, 2 mM Na_2_ATP, and 0.3 mM Na_2_GTP. AMPA receptor-mediated currents were blocked by adding 10 μM NBQX to the aCSF. A stable gigaseal was formed between the electrode and PC membrane, followed by negative pressure to rupture the membrane and establish whole-cell configuration (access resistance < 20 MΩ, leak current < −30 pA). Membranes were voltage-clamped at −70 mV after stabilization.

Spontaneous action potentials (sAP) were recorded in cell-attached mode. Negative pressure was applied to gradually increase the resistance between the electrode and the cell membrane to 200 to 300 MΩ. After a brief stabilization period, cell-attached recording was initiated.

Electrophysiological data were acquired using Clampex 10.7 software and analyzed using Clampfit 10.7 software.

### Single cell RNA sequencing

ECs were isolated from the cerebellum of *Prmt5^fl/fl^;ROSA26^LSL-tdTomato^* mice and *Prmt5^fl/+^;ROSA26^LSL-tdTomato^*littermates at P20 using fluorescence-activated cell sorting (FACS). Briefly, ECs were stained with APC rat anti-mouse CD31 (eBiosciences, 17-0311-82) and PE rat anti-mouse CD45 (eBiosciences, 12-0453-82) for 1 hour at 4°C. Cell viability was assessed using 7-AAD (eBiosciences, 00-6993-50) to exclude dead cells. Viable ECs (RFP^+^CD31^+^CD45^-^7-AAD^-^) were sorted using a BD FACSAria III sorter (BD Biosciences). Freshly sorted cells were immediately processed for droplet-based single-cell RNA sequencing (10x Genomics Chromium platform). Single-cell libraries were prepared using the Chromium Single Cell 3ʹ Reagent Kits v3 (10x Genomics) following the manufacturer’s protocol. Briefly, FACS-sorted cells were washed three times with Dulbecco’s phosphate-buffered saline (DPBS) containing 0.04% BSA and resuspended at 700-1200 cells/µL, with viability of ≥ 85%. Cellular encapsulation was performed using the Chromium Controller (10x Genomics), with cell capture efficiency monitored in real-time. Reverse transcription within droplets was conducted to generate uniquely barcoded cDNA. Emulsions were disrupted post-reverse transcription, and barcoded cDNA was purified using Dynabeads. The purified cDNA was amplified by PCR. For 3ʹ gene expression library construction, 50 ng of amplified cDNA was fragmented, end-repaired, and subjected to double size selection using SPRIselect beads. Final libraries were sequenced on an Illumina NovaSeq X platform to generate 150 bp paired-end reads.

For 10x Genomic data processing, raw reads were demultiplexed and mapped to the reference genome by 10X Genomics Cell Ranger (v1.1.0) with default parameters. To ensure data quality, we meticulously filtered the cells, retaining only those that met the following criteria: (1) Cell-level filtering (300 < detected genes < 5000; total UMI counts > 1000; exclusion of the top 3% high-UMI outliers) (2) Hemoglobin gene filtering (Hb[%] < 0.5) (3) Mitochondrial content thresholding (MT[%] < 10). Data normalization and clustering were performed in Seurat (v5.2.1) using SCTransform normalization. For graph-based clustering, we selected 2000 highly variable genes (HVGs) and performed principal component analysis (PCA) on scaled data. A shared nearest neighbor (SNN) graph was constructed using the first 30 principal components and clustered with Leiden algorithm (resolution=0.5). Cell population clusters were annotated to known cell types by evaluating the expression of specific marker genes previously identified in the literature. Differential expression analysis employed the Wilcoxon rank-sum test, and only genes with FDR < 0.05 and |log2FC| > 0.5) were considered to be DEGs. Functional enrichment of DEGs was analyzed using ClusterProfiler (v4.14.6) with biological process terms from the Gene Ontology (GO) database.

### Statistical Analysis

Detailed statistical information for all experiments, including the specific tests applied, exact sample sizes (n), and the definition of n (e.g., number of biological replicates, cells, or independent experiments), is provided in the respective figure legends. Unless otherwise stated, data are presented as mean ± SEM. Comparisons between two groups were analyzed using an unpaired two-tailed Student’s t-test. Statistical significance was defined as P < 0.05. All statistical analyses were performed with GraphPad Prism software.

## Acknowledgements

We would like to thank Dr. Haitao Wu and Dr. Zhiying Wu for helpful discussions relating to phenotype analysis. We are further indebted to Dr. Yichang Jia, Dr. Haitao Li, and Dr. Zhijie Chang for their rigorous critique and constructive recommendations pertaining to this study.

## Author contributions

Conceptualization: X.Y. and J.W. Formal analysis: J.L. and Y.Z. Investigation: J.L., Y.Z., Y.H., H.N., Z.F., S.H., Y.C., Y.H., Y.C., W.L., H.A., and S.X. Methodology: J.L. and Y.Z. Project administration: X.Y. and J.W. Supervision: X.Y. and J.W. Writing—original draft: X.Y., W.J., J.L., Y.Z. and Y.H. Writing—review & editing: All authors.

## Data and materials availability

All data are available in the main text or the supplementary materials. Any additional information necessary to reanalyze the data presented in this study is available from the Lead Contact upon reasonable request.

## References

1. S. Schaeffer, C. Iadecola, Revisiting the neurovascular unit. Nat. Neurosci. 24, 1198–1209 (2021).

2. M. Segarra, M. R. Aburto, J. Hefendehl, A. Acker-Palmer, Neurovascular interactions in the nervous system. Annu. Rev. Cell Dev. Biol. 35, 615–635 (2019).

3. I. Paredes, P. Himmels, C. Ruiz de Almodóvar, Neurovascular communication during CNS development. Dev. Cell 45, 10–32 (2018).

4. S. Li, T. P. Kumar, S. Joshee, T. Kirschstein, S. Subburaju, J. S. Khalili, J. Kloepper, C. Du, A. Elkhal, G. Szabó, R. K. Jain, R. Köhling, A. Vasudevan, Endothelial cell-derived GABA signaling modulates neuronal migration and postnatal behavior. Cell Res. 28, 221–248 (2018).

5. C. Tan, N. N. Lu, C. K. Wang, D. Y. Chen, N. H. Sun, H. Lyu, J. Körbelin, W. X. Shi, K. Fukunaga, Y. M. Lu, F. Han, Endothelium-derived Semaphorin 3G regulates hippocampal synaptic structure and plasticity via Neuropilin-2/PlexinA4. Neuron 101, 920–937.e13 (2019).

6. A. C. Delgado, S. R. Ferrón, D. Vicente, E. Porlan, A. Perez-Villalba, C. M. Trujillo, P. D’Ocón, I. Fariñas, Endothelial NT-3 delivered by vasculature and CSF promotes quiescence of subependymal neural stem cells through nitric oxide induction. Neuron 83, 572–585 (2014).

7. M. Zhang, L. Su, W. Wang, C. Li, Q. Liang, F. Ji, J. Jiao, Endothelial cells regulated by RNF20 orchestrate the proliferation and differentiation of neural precursor cells during embryonic development. Cell Rep. 40, 111350 (2022).

8. K. W. Wu, L. L. Lv, Y. Lei, C. Qian, F. Y. Sun, Endothelial cells promote excitatory synaptogenesis and improve ischemia-induced motor deficits in neonatal mice. Neurobiol. Dis. 121, 230–239 (2019).

9. J. Wang, Y. Cui, Z. Yu, W. Wang, X. Cheng, W. Ji, S. Guo, Q. Zhou, N. Wu, Y. Chen, Y. Chen, X. Song, H. Jiang, Y. Wang, Y. Lan, B. Zhou, L. Mao, J. Li, H. Yang, W. Guo, X. Yang, Brain endothelial cells maintain lactate homeostasis and control adult hippocampal neurogenesis. Cell Stem Cell 25, 754–767.e9 (2019).

10. L. N. Nguyen, D. Ma, G. Shui, P. Wong, A. Cazenave-Gassiot, X. Zhang, M. R. Wenk, E. L. Goh, D. L. Silver, Mfsd2a is a transporter for the essential omega-3 fatty acid docosahexaenoic acid. Nature 509, 503–506 (2014).

11. Y. Chen, J. Joo, J. M. Chu, R. C. Chang, G. T. Wong, Downregulation of the glucose transporter GLUT 1 in the cerebral microvasculature contributes to postoperative neurocognitive disorders in aged mice. J. Neuroinflammation 20, 237 (2023).

12. S. Mayerl, J. Müller, R. Bauer, S. Richert, C. M. Kassmann, V. M. Darras, K. Buder, A. Boelen, T. J. Visser, H. Heuer, Transporters MCT8 and OATP1C1 maintain murine brain thyroid hormone homeostasis. J. Clin. Invest. 124, 1987–1999 (2014).

13. M. D. Sweeney, Z. Zhao, A. Montagne, A. R. Nelson, B. V. Zlokovic, Blood-brain barrier: From physiology to disease and back. Physiol. Rev. 99, 21–78 (2019).

14. C. I. De Zeeuw, S. G. Lisberger, J. L. Raymond, Diversity and dynamism in the cerebellum. Nat. Neurosci. 24, 160–167 (2021).

15. C. Hull, W. G. Regehr, The cerebellar cortex. Annu. Rev. Neurosci. 45, 151–175 (2022).

16. V. Kozareva, C. Martin, T. Osorno, S. Rudolph, C. Guo, C. Vanderburg, N. Nadaf, A. Regev, W. G. Regehr, E. Macosko, A transcriptomic atlas of mouse cerebellar cortex comprehensively defines cell types. Nature 598, 214–219 (2021).

17. D. V. Jeste, L. Barban, J. Parisi, Reduced Purkinje cell density in Huntington’s disease. Exp. Neurol. 85, 78–86 (1984).

18. E. D. Louis, P. L. Faust, J. P. Vonsattel, L. S. Honig, A. Rajput, C. A. Robinson, A. Rajput, R. Pahwa, K. E. Lyons, G. W. Ross, S. Borden, C. B. Moskowitz, A. Lawton, N. Hernandez, Neuropathological changes in essential tremor: 33 cases compared with 21 controls. Brain 130, 3297–3307 (2007).

19. M. Salcman, R. Defendini, J. Correll, S. Gilman, Neuropathological changes in cerebellar biopsies of epileptic patients. Ann. Neurol. 3, 10–19 (1978).

20. T. Klockgether, C. Mariotti, H. L. Paulson, Spinocerebellar ataxia. Nat. Rev. Dis. Primers 5, 24 (2019).

21. F. Pilotto, C. Douthwaite, R. Diab, X. Ye, Z. Al Qassab, C. Tietje, M. Mounassir, A. Odriozola, A. Thapa, R. A. M. Buijsen, S. Lagache, A. C. Uldry, M. Heller, S. Müller, W. M. C. van Roon-Mom, B. Zuber, S. Liebscher, S. Saxena, Early molecular layer interneuron hyperactivity triggers Purkinje neuron degeneration in SCA1. Neuron 111, 2523–2543.e10 (2023).

22. C. C. Lin, K. C. Fang, I. Balbo, T. Y. Liang, C. W. Liu, W. C. Liu, Y. M. Wang, Y. L. Hung, K. C. Yang, S. K. Geng, C. L. Ni, C. P. Driscoll, D. S. Ruff, A. Kumar, N. Amokrane, N. Desai, P. L. Faust, E. D. Louis, S. H. Kuo, M. K. Pan, Reduced cerebellar rhythm by climbing fiber denervation is linked to motor rhythm deficits in mice and ataxia severity in patients. Sci. Transl. Med. 17, eadk3922 (2025).

23. M. Tolve, J. Tutas, E. Özer-Yildiz, I. Klein, A. Petzold, V. J. Fritz, M. Overhoff, Q. Silverman, E. Koletsou, F. Liebsch, G. Schwarz, T. Korotkova, S. Valtcheva, G. Gatto, N. L. Kononenko, The endocytic adaptor AP-2 maintains Purkinje cell function by balancing cerebellar parallel and climbing fiber synapses. Cell Rep 44, 115256 (2025).

24. K. J. Robinson, M. Watchon, A. S. Laird, Aberrant cerebellar circuitry in the spinocerebellar ataxias. Front. Neurosci. 14, 707 (2020).

25. H. Takahashi, K. Katayama, K. Sohya, H. Miyamoto, T. Prasad, Y. Matsumoto, M. Ota, H. Yasuda, T. Tsumoto, J. Aruga, A. M. Craig, Selective control of inhibitory synapse development by Slitrk3-PTPδ trans-synaptic interaction. Nat. Neurosci. 15, 389–398, s381-382 (2012).

26. J. Woo, S. K. Kwon, J. Nam, S. Choi, H. Takahashi, D. Krueger, J. Park, Y. Lee, J. Y. Bae, D. Lee, J. Ko, H. Kim, M. H. Kim, Y. C. Bae, S. Chang, A. M. Craig, E. Kim, The adhesion protein IgSF9b is coupled to neuroligin 2 via S-SCAM to promote inhibitory synapse development. J. Cell Biol. 201, 929–944 (2013).

27. K. Lee, Y. Kim, S. J. Lee, Y. Qiang, D. Lee, H. W. Lee, H. Kim, H. S. Je, T. C. Südhof, J. Ko, MDGAs interact selectively with neuroligin-2 but not other neuroligins to regulate inhibitory synapse development. Proc. Natl. Acad. Sci. U.S. A. 110, 336–341 (2013).

27. S. Früh, J. Romanos, P. Panzanelli, D. Bürgisser, S. K. Tyagarajan, K. P. Campbell, M. Santello, J. M. Fritschy, Neuronal dystroglycan is necessary for formation and maintenance of functional CCK-positive basket cell terminals on pyramidal cells. J. Neurosci. 36, 10296–10313 (2016).

28. F. Varoqueaux, S. Jamain, N. Brose, Neuroligin 2 is exclusively localized to inhibitory synapses. Eur. J. Cell Biol. 83, 449–456 (2004).

29. D. Irala, S. Wang, K. Sakers, L. Nagendren, F. P. Ulloa Severino, D. S. Bindu, J. T. Savage, C. Eroglu, Astrocyte-secreted neurocan controls inhibitory synapse formation and function. Neuron 112, 1657–1675.e10 (2024).

30. A. Terauchi, E. M. Johnson-Venkatesh, A. B. Toth, D. Javed, M. A. Sutton, H. Umemori, Distinct FGFs promote differentiation of excitatory and inhibitory synapses. Nature 465, 783–787 (2010).

31. M. Yasumura, T. Yoshida, S. J. Lee, T. Uemura, J. Y. Joo, M. Mishina, Glutamate receptor δ1 induces preferentially inhibitory presynaptic differentiation of cortical neurons by interacting with neurexins through cerebellin precursor protein subtypes. J. Neurochem. 121, 705–716 (2012).

32. M. Yuzaki, Two classes of secreted synaptic organizers in the central nervous system. Annu. Rev. Physiol. 80, 243–262 (2018).

33. Q. Wu, M. Schapira, C. H. Arrowsmith, D. Barsyte-Lovejoy, Protein arginine methylation: from enigmatic functions to therapeutic targeting. Nat. Rev. Drug Discov. 20, 509–530 (2021).

34. Q. Ye, J. Zhang, C. Zhang, B. Yi, K. Kazama, W. Liu, X. Sun, Y. Liu, J. Sun, Endothelial PRMT5 plays a crucial role in angiogenesis after acute ischemic injury. JCI Insight 7, e152481 (2022).

35. A. Quillien, G. Gilbert, M. Boulet, S. Ethuin, L. Waltzer, L. Vandel, Prmt5 promotes vascular morphogenesis independently of its methyltransferase activity. PLoS Genet. 17, e1009641 (2021).

36. Z. Li, J. Xu, Y. Song, C. Xin, L. Liu, N. Hou, Y. Teng, X. Cheng, T. Wang, Z. Yu, J. Song, Y. Zhang, J. Wang, X. Yang, PRMT5 prevents dilated cardiomyopathy via suppression of protein O-GlcNAcylation. Circ. Res. 129, 857–871 (2021).

37. F. Li, Y. Lan, Y. Wang, J. Wang, G. Yang, F. Meng, H. Han, A. Meng, Y. Wang, X. Yang, Endothelial Smad4 maintains cerebrovascular integrity by activating N-cadherin through cooperation with Notch. Dev. Cell 20, 291–302 (2011).

38. F. Meng, L. Shi, X. Cheng, N. Hou, Y. Wang, Y. Teng, A. Meng, X. Yang, Surfactant protein A promoter directs the expression of Cre recombinase in brain microvascular endothelial cells of transgenic mice. Matrix Biol. 26, 54–57 (2007).

39. F. Côté, J. F. Collard, J. P. Julien, Progressive neuronopathy in transgenic mice expressing the human neurofilament heavy gene: a mouse model of amyotrophic lateral sclerosis. Cell 73, 35–46 (1993).

40. L. Zhang, J. Zhou, W. Kong, Extracellular matrix in vascular homeostasis and disease. Nat. Rev. Cardiol. 22, 333–353 (2025).

41. A. Naba, Mechanisms of assembly and remodelling of the extracellular matrix. Nat. Rev. Mol. Cell Biol. 25, 865–885 (2024).

42. E. S. Novoseletskaya, P. V. Evdokimov, A. Y. Efimenko, Extracellular matrix-induced signaling pathways in mesenchymal stem/stromal cells. Cell Commun. Signal. 21, 244 (2023).

43. G. E. Davis, S. S. Kemp, Extracellular matrix regulation of vascular morphogenesis, maturation, and stabilization. Cold Spring Harb. Perspect. Med. 13, a041156 (2023).

44. M. H. Yau, Y. Wang, K. S. Lam, J. Zhang, D. Wu, A. Xu, A highly conserved motif within the NH2-terminal coiled-coil domain of angiopoietin-like protein 4 confers its inhibitory effects on lipoprotein lipase by disrupting the enzyme dimerization. J. Biol. Chem. 284, 11942–11952 (2009).

45. B. Chaube, K. M. Citrin, M. Sahraei, A. K. Singh, D. S. de Urturi, W. Ding, R. W. Pierce, R. Raaisa, R. Cardone, R. Kibbey, C. Fernández-Hernando, Y. Suárez, Suppression of angiopoietin-like 4 reprograms endothelial cell metabolism and inhibits angiogenesis. Nat. Commun. 14, 8251 (2023).

46. R. L. Huang, Z. Teo, H. C. Chong, P. Zhu, M. J. Tan, C. K. Tan, C. R. Lam, M. K. Sng, D. T. Leong, S. M. Tan, S. Kersten, J. L. Ding, H. Y. Li, N. S. Tan, ANGPTL4 modulates vascular junction integrity by integrin signaling and disruption of intercellular VE-cadherin and claudin-5 clusters. Blood 118, 3990–4002 (2011).

47. X. Zheng, S. Shen, A. Wang, Z. Zhu, Y. Peng, H. Peng, C. Zhong, D. Guo, T. Xu, J. Chen, Z. Ju, D. Geng, Y. Zhang, J. He, Angiopoietin-like protein 4 and clinical outcomes in ischemic stroke patients. Ann. Clin. Transl. Neurol. 8, 687–695 (2021).

48. M. Ruscica, F. Zimetti, M. P. Adorni, C. R. Sirtori, M. G. Lupo, N. Ferri, Pharmacological aspects of ANGPTL3 and ANGPTL4 inhibitors: New therapeutic approaches for the treatment of atherogenic dyslipidemia. Pharmacol. Res. 153, 104653 (2020).

49. Q. Zhao, G. Rank, Y. T. Tan, H. Li, R. L. Moritz, R. J. Simpson, L. Cerruti, D. J. Curtis, D. J. Patel, C. D. Allis, J. M. Cunningham, S. M. Jane, PRMT5-mediated methylation of histone H4R3 recruits DNMT3A, coupling histone and DNA methylation in gene silencing. Nat. Struct. Mol. Biol. 16, 304–311 (2009).

50. S. Pal, S. N. Vishwanath, H. Erdjument-Bromage, P. Tempst, S. Sif, Human SWI/SNF-associated PRMT5 methylates histone H3 arginine 8 and negatively regulates expression of ST7 and NM23 tumor suppressor genes. Mol. Cell. Biol. 24, 9630–9645 (2004).

51. Y. C. Chen, E. Koutelou, S. Y. R. Dent, Now open: Evolving insights to the roles of lysine acetylation in chromatin organization and function. Mol. Cell 82, 716–727 (2022).

52. F. J. Garcia, M. Heiman, Molecular and cellular characteristics of cerebrovascular cell types and their contribution to neurodegenerative diseases. Mol. Neurodegener. 20, 13 (2025).

53. M. T. Jeon, K. S. Kim, E. S. Kim, S. Lee, J. Kim, H. S. Hoe, D. G. Kim, Emerging pathogenic role of peripheral blood factors following BBB disruption in neurodegenerative disease. Ageing Res. Rev. 68, 101333 (2021).

54. C. Bouleti, T. Mathivet, B. Coqueran, J. M. Serfaty, M. Lesage, E. Berland, C. Ardidie-Robouant, G. Kauffenstein, D. Henrion, B. Lapergue, M. Mazighi, C. Duyckaerts, G. Thurston, D. M. Valenzuela, A. J. Murphy, G. D. Yancopoulos, C. Monnot, I. Margaill, S. Germain, Protective effects of angiopoietin-like 4 on cerebrovascular and functional damages in ischaemic stroke. Eur. Heart J. 34, 3657–3668 (2013).

55. Z. Qiu, J. Yang, G. Deng, D. Li, S. Zhang, Angiopoietin-like 4 promotes angiogenesis and neurogenesis in a mouse model of acute ischemic stroke. Brain Res. Bull. 168, 156–164 (2021).

56. H. Li, J. Wei, Z. Zheng, R. Wang, M. Qu, J. Liu, G. Lu, X. Li, W. Gong, The therapeutic potential of recombinant ANGPTL4 in Parkinson’s disease: Evidence from in vivo and in vitro studies. Free Radic. Biol. Med. 227, 190–200 (2025).

57. A. Chakraborty, A. Kamermans, B. van Het Hof, K. Castricum, E. Aanhane, J. van Horssen, V. L. Thijssen, P. Scheltens, C. E. Teunissen, R. D. Fontijn, W. M. van der Flier, H. E. de Vries, Angiopoietin like-4 as a novel vascular mediator in capillary cerebral amyloid angiopathy. Brain 141, 3377–3388 (2018).

58. L. Liu, J. Li, D. Huo, Z. Peng, R. Yang, J. Fu, B. Xu, B. Yang, H. Chen, X. Wang, Meningitic escherichia coli induction of ANGPTL4 in brain microvascular endothelial cells contributes to blood-brain barrier disruption via ARHGAP5/RhoA/MYL5 signaling cascade. Pathogens 8, 254 (2019).

59. C. Yao, J. Wang, ANGPTL4 promoted the cognitive impairment in vascular dementia via increasing integrin/p-Syk signalings induced mitochondrial autophagy and apoptosis in the hippocampus. Sci. Rep. 15, 25312 (2025).

60. N. Li, X. Wang, R. Lin, F. Yang, H. C. Chang, X. Gu, J. Shu, G. Liu, Y. Yu, W. Wei, Z. Bao, ANGPTL4-mediated microglial lipid droplet accumulation: Bridging Alzheimer’s disease and obesity. Neurobiol. Dis. 203, 106741 (2024).

61. F. Xiong, X. Liao, J. Xiao, X. Bai, J. Huang, B. Zhang, F. Li, P. Li, Emerging limb rehabilitation therapy after post-stroke motor recovery. Front. Aging Neurosci. 14, 863379 (2022).

62. C. Su, X. Yang, S. Wei, R. Zhao, Association of cerebral small vessel disease with gait and balance disorders. Front. Aging Neurosci. 14, 834496 (2022).

