## Supplementary Table for "Brain endothelial PRMT5-ANGPTL4 axis regulates cerebellar inhibitory synaptogenesis and motor coordination"

| Supplementary Table 1 | | | | | | |
|---|---|---|---|---|---|---|
| **Name** | **Source** | **Catalog Number** | **Dilution** | **Reactivity** | **Second antibody** | **Remark** |
| anti-VGAT | Proteintech | Cat# 14471-1-AP; RRID: AB_10644324 | 1:500 | Mouse | Rabbit | IF-P/WB |
| anti-VGAT | GeneTex | Cat# GTX101908; RRID: AB_10619521 | 1:500 | Mouse | Rabbit | ICC |
| anti-VGLUT1 | Abcam | Cat# ab227805; RRID: AB_2868428 | 1:500 | Mouse | Rabbit | |
| anti-VGLUT2 | Abcam | Cat# ab216463; RRID: AB_2893024 | 1:500 | Mouse | Rabbit | |
| anti-Calbindin | Cell Signaling Technology | Cat# 13176; RRID: AB_2687400 | 1:500 | Mouse | Rabbit | |
| anti-MAP2 | GeneTex | Cat# GTX634473; RRID: AB_2888458 | 1:500 | Mouse | Mouse | |
| anti-CD31 | Abcam | Cat# ab28364; RRID: AB_726362 | 1:500 | Mouse | Rabbit | IF-P |
| anti-CD31 | BD Bioscience | Cat# 550274; RRID: AB_393571 | 1:200 | Mouse | Rat | IF-Fr |
| anti-NeuN | Abcam | Cat# ab177487; RRID: AB_2532109 | 1:500 | Mouse | Rabbit | |
| anti-GFAP | Cell Signaling Technology | Cat# 3670; RRID: AB_561049 | 1:500 | Mouse | Mouse | |
| anti-Iba1 | Abcam | Cat# ab178846; RRID: AB_2636859 | 1:500 | Mouse | Rabbit | |
| anti-PRMT5 | MedChemExpress | Cat# HY-P82108; RRID: AB_3675821 | 1:500 | Mouse | Rabbit | IF-P/WB |
| anti-ERG | Abcam | Cat# ab92513; RRID: AB_2630401 | 1:500 | Mouse | Rabbit | |
| anti-ANGPTL4 | Proteintech | Cat# 67577-1-IG; RRID: AB_2882790 | 1:1000 | Mouse | Mouse | |
| anti-GAPDH | ZSGB-BIO | Cat# TA-08; RRID: AB_2747414 | 1:2000 | Mouse | Mouse | |
| Biotin anti-mouse CD31 Antibody | BioLegend | Cat# 102504; RRID: AB_312910 | | | | |
| HRP Goat Anti-Rabbit IgG Polymer Detection Kit | Vector Laboratories | Cat# MP-7451; RRID: AB_2631198 | | | | |
| Goat anti-mouse IgG | ZSGB-BIO | Cat# ZB-2305; RRID: AB_2747415 | | | | |

| | | | | | | |
|---|---|---|---|---|---|---|
| Goat anti-Rabbit IgG | ZSGB-BIO | Cat# ZB-2301; RRID: AB_2747412 | | | | |
| Rabbit IgG antibody | Abcam | Cat# ab171870; RRID: AB_2687657 | | | | ChIP |
| PRMT5 antibody | GeneTex | Cat# GTX116004; RRID: AB_10624553 | | | | ChIP |
| Histone H4R3me2s (symmetric) antibody | Active Motif | Cat# 61187; RRID: AB_2793544 | | | | |
| Histone H3R8 Dimethyl Symmetric antibody | Epigentek | Cat# A-3706; RRID: AB_2620161 | | | | |
| Histone H3K9ac antibody | Active Motif | Cat# 39038; RRID: AB_2561017 | | | | |

| Supplementary Table 2 | | |
|---|---|---|
| **Name** | **Source** | **Catalog Number** |
| StemPro$^{TM}$ Accutase$^{TM}$ Cell Dissociation Reagent | Gibco | Cat# A1110501 |
| GlutaMAX Supplement | Gibco | Cat# 35050-061 |
| B27 Supplement | Gibco | Cat# 17504-044 |
| Fetal Bovine Serum (FBS) | VivaCell | Cat# C04001-500 |
| DMEM | VivaCell | Cat# C3113-0500 |
| Neurobasal™ Medium | Gibco | Cat# 21103049 |
| Endothelial Cell Growth Medium-2 (EGM-2) | Lonza Bioscience | Cat# CC-3162 |
| Penicillin-Streptomycin | VivaCell | Cat# C3420-0100 |
| TRIzol | Invitrogen | Cat# 15596026 |
| EZ-Link$^{TM}$ Sulfo-NHS-LC-Biotin | Thermo Fisher | Cat# 21335 |
| jetPRIME® | Polyplus | Cat# 101000046 |
| DAPI | Invitrogen | Cat# D1306 |
| Goat serum | ZSGB-BIO | Cat# ZLI-9056 |
| RIPA lysis buffer | Applygen | Cat# C1053 |
| Angiopoietin-related protein 4/ANGPTL4 Protein | MedChemExpress | Cat# HY-P7508 |
| DNase I | Roche | Cat# 10104159001 |
| Collagenase Type I | Sigma | Cat# C0130 |
| Bovine Serum Albumin | Sigma | Cat# V900933 |
| Enlight$^{TM}$ Western blot substrate | Engreen Biosystem | Cat# 29100 |
| BCA Protein Assay Reagent | Thermo Fisher | Cat# 23225 |
| TSA-fluorophores Working Solution | Histova Biotechnology | Cat# NECC7100 |

| | | |
|---|---|---|
| Mouse Angiopoietin-related protein 4 (ANGPTL4) ELISA kit | CUSABIO | Cat# CSB-EL001712MO |
| Reverse transcription kit | TOYOBO | Cat# FSQ-201 |
| SYBR Green Real-time PCR Master Mix | TOYOBO | Cat# QPK-201 |
