## Supplementary material for "Brain endothelial PRMT5-ANGPTL4 axis regulates cerebellar inhibitory synaptogenesis and motor coordination": Figure S1-S8

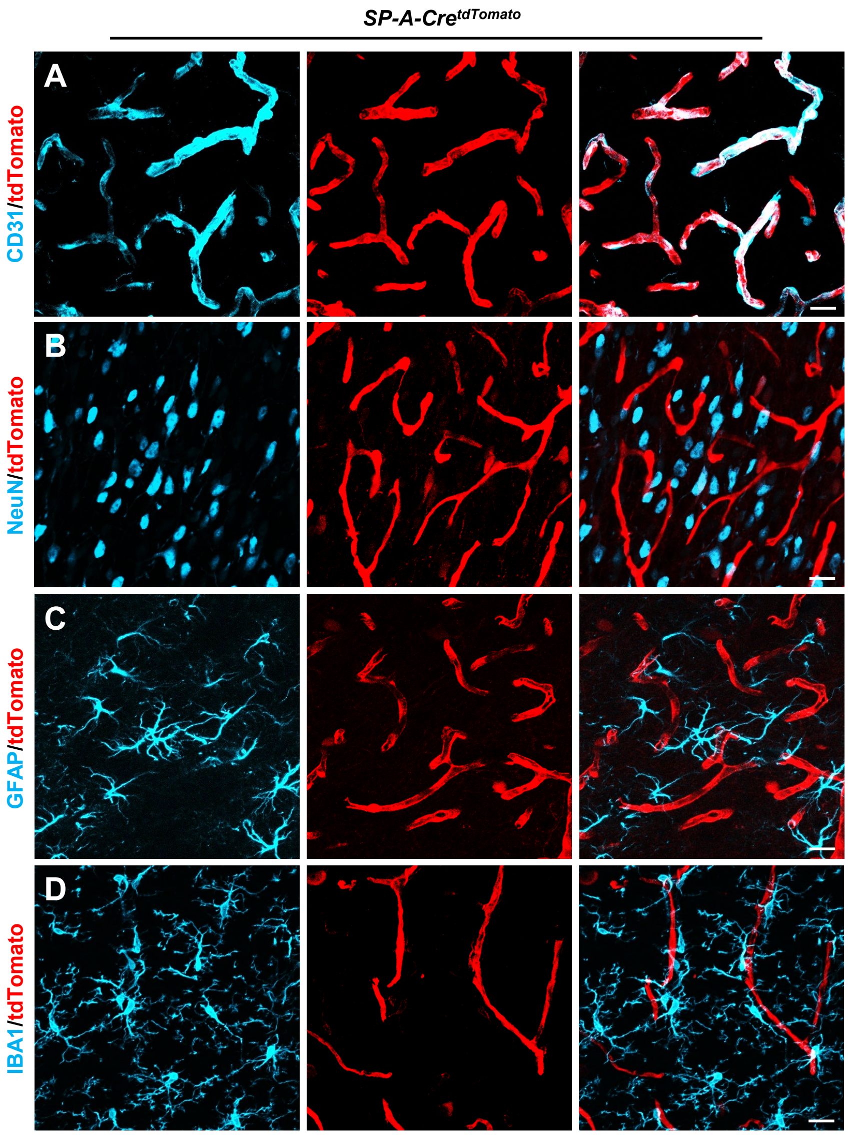


**Fig. S1. *SP-A-Cre* is specifically expressed in brain ECs**

(A-D) Confocal images of tdTomato (red), CD31 (blue), NeuN (blue), GFAP (blue), and IBA1 (blue) immunostaining in P60 *SP-A-Cre*$^{tdTomato}$ mouse brain sections. Scale bar, 20 μm.

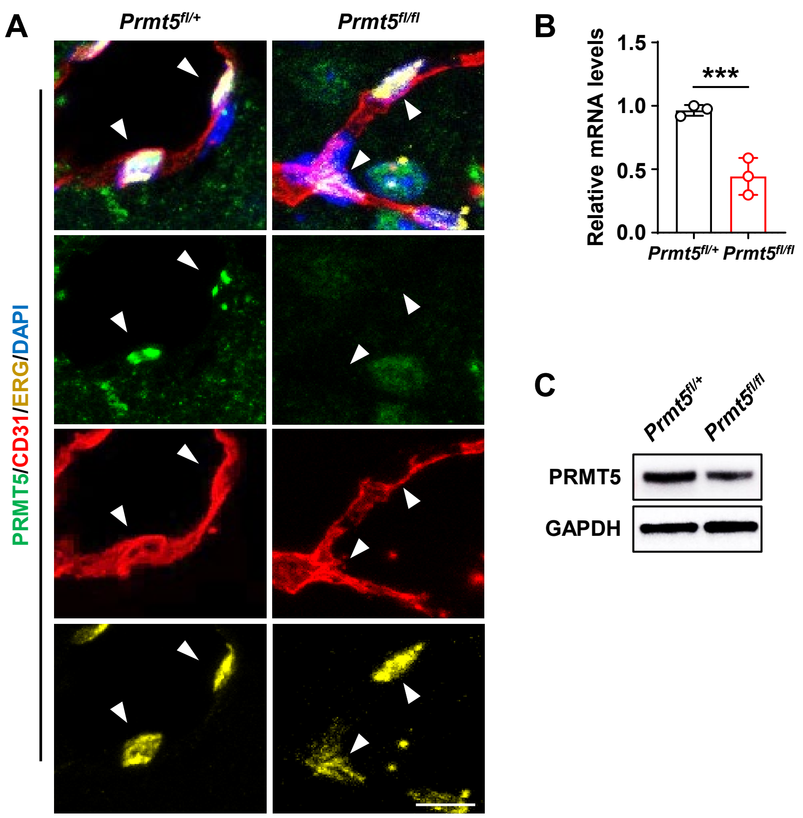


**Fig. S2. Reduced PRMT5 expression in brain ECs of *Prmt5$^{fl/fl}$* mice**

(A) Confocal images of PRMT5 (green), CD31 (red), and ERG (yellow) immunostaining in brain sections from P60 *Prmt5$^{fl/+}$* and *Prmt5$^{fl/fl}$* mice. Scale bar, 10 µm.

(B) qPCR analysis of *Prmt5* mRNA levels in brain ECs isolated from P20 *Prmt5$^{fl/+}$* and *Prmt5$^{fl/fl}$* mice. ***P < 0.001 (mean ± SEM, n = 3 per group).

(C) Western blot analysis of PRMT5 expression in brain ECs from P20 *Prmt5$^{fl/+}$* and *Prmt5$^{fl/fl}$* mice. GAPDH served as the loading control.

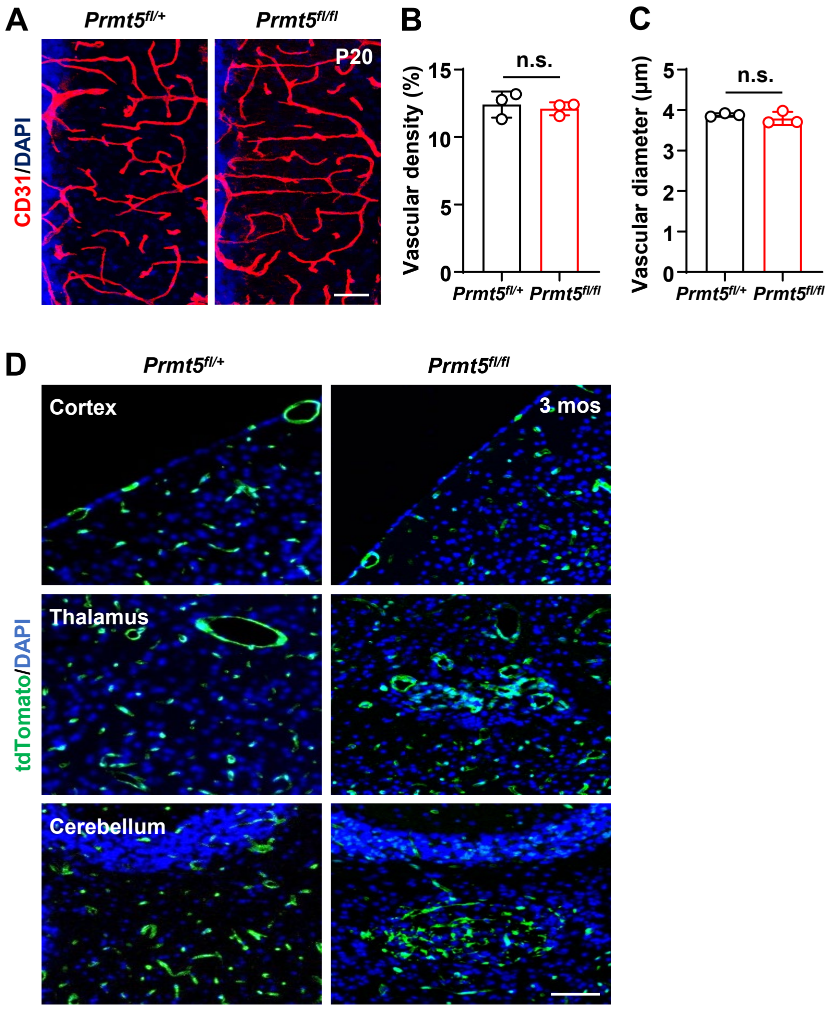


**Fig. S3. Brain endothelial deletion of *Prmt5* induces progressive cerebrovascular anomalies**

(A) Confocal images of CD31 (red) immunostaining showing cerebrovascular morphology in P20 *Prmt5*$^{fl/+}$ and *Prmt5*$^{fl/fl}$ mice. Scale bar, 50 μm.

(B and C) Bar graphs showing cerebrovascular density (B) and diameter (C) in P20 *Prmt5*$^{fl/+}$ and *Prmt5*$^{fl/fl}$ mice. n.s., not significant (mean ± SEM, n = 3 mice per group).

(D) Confocal images of tdTomato (green) immunostaining showing cerebrovascular malformation in the cerebellar, thalamic, and cortical regions of 3-month-old *Prmt5*$^{fl/+}$ and *Prmt5*$^{fl/fl}$ mice. Scale bar, 50 μm.

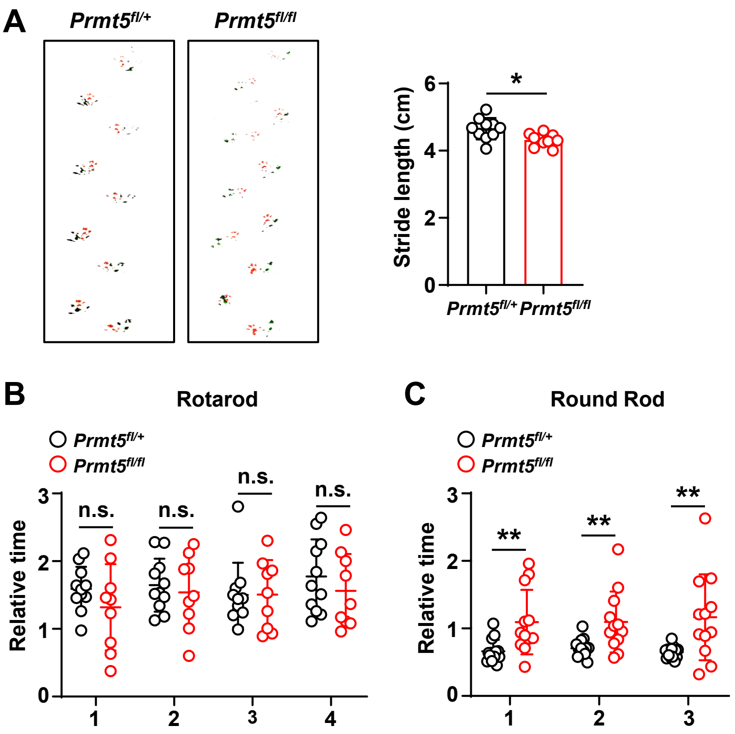


**Fig. S4. Loss of endothelial *Prmt5* leads to mild motor deficits in *Prmt5*$^{fl/fl}$ mice at P20**

(A) Representative footprints and quantification of stride length in *Prmt5*$^{fl/+}$ and *Prmt5*$^{fl/fl}$ mice. Red, forepaws; green, hindpaws. *P < 0.05 (mean ± SEM, n = 10 mice per group).

(B) Quantification of latency time spent on the rotarod. Data are from four independent trials. n.s., not significant (mean ± SEM, n = 11 mice in *Prmt5*$^{fl/+}$ group, n = 9 mice in *Prmt5*$^{fl/fl}$ group).

(C) Quantification of time spent crossing the balance beam. Data are from three independent trials. **P < 0.01 (mean ± SEM, n = 14 mice in *Prmt5*$^{fl/+}$ group, n = 12 mice in *Prmt5*$^{fl/fl}$ group).

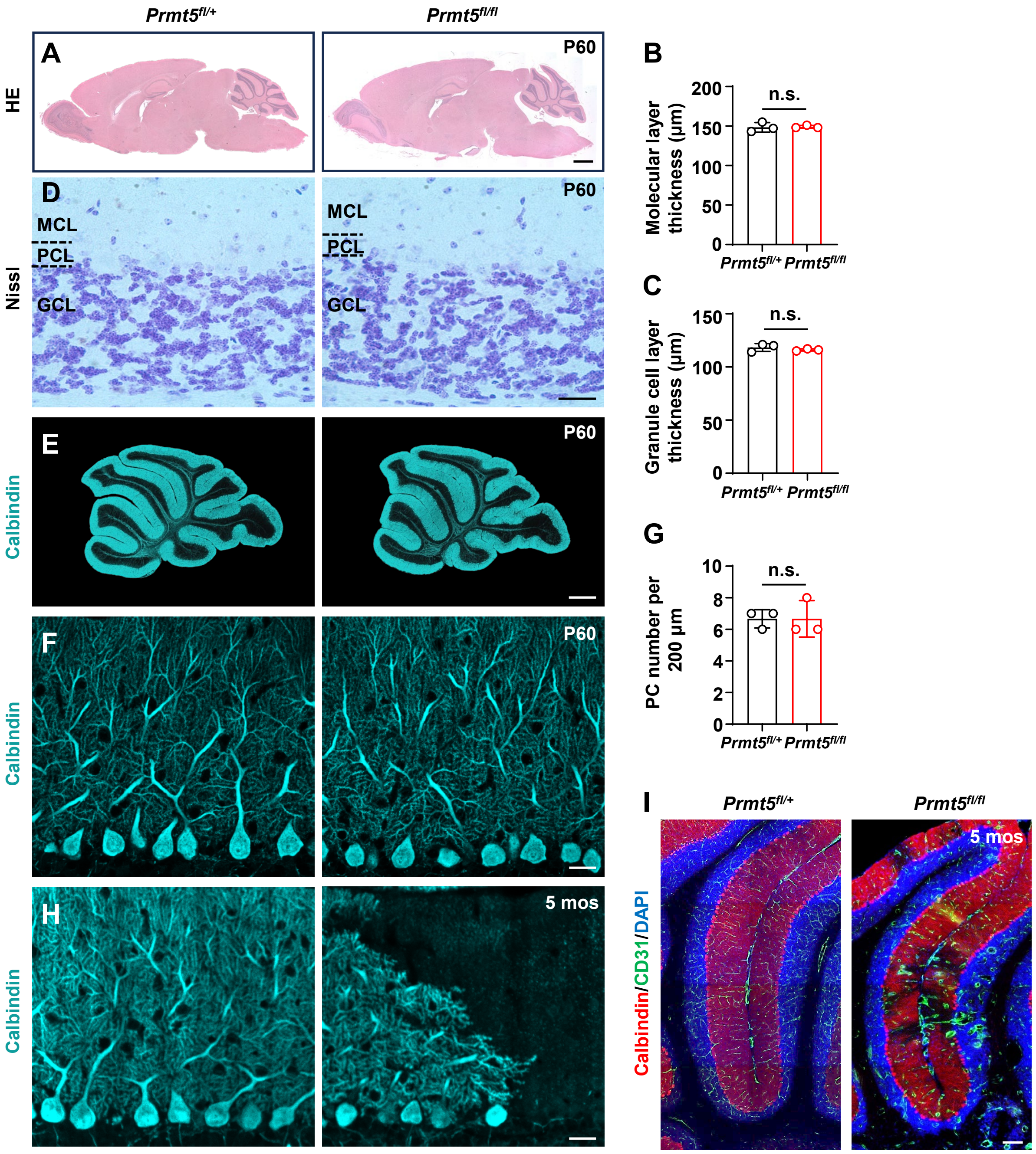


**Fig. S5. *Prmt5*$^{fl/fl}$ mice maintain normal cerebellar structure at P60, followed by progressive PC degeneration at 5 months**

(A) HE staining of sagittal brain sections from P60 *Prmt5*$^{fl/+}$ and *Prmt5*$^{fl/fl}$ mice. Scale bar, 1,000 μm.

(B and C) Bar plots showing molecular layer thickness (B) and granule cell layer thickness (C). n.s., not significant (mean ± SEM, n = 3 mice per group).

(D) Nissl staining of sagittal cerebellar cortex sections from P60 *Prmt5*$^{fl/+}$ and *Prmt5*$^{fl/fl}$ mice. Scale bar, 25μm.

(E) Confocal images of Calbindin (blue) immunostaining in P60 *Prmt5*$^{fl/+}$ and *Prmt5*$^{fl/fl}$ mice. Scale bar, 500 μm.

(F) Confocal images of Calbindin (blue) immunostaining in P60 *Prmt5*$^{fl/+}$ and *Prmt5*$^{fl/fl}$ mice. Scale bar, 20 μm.

(G) Bar plots showing PC number. n.s., not significant (mean ± SEM, n = 3 mice per group).

(H) Confocal images of Calbindin (blue) immunostaining in P60 and 5-month-old *Prmt5*$^{fl/+}$ and *Prmt5*$^{fl/fl}$ mice. Scale bar, 20 μm.

(I) Confocal images of Calbindin (red) and CD31 (green) immunostaining in 5-month-old *Prmt5*$^{fl/+}$ and *Prmt5*$^{fl/fl}$ mice. Scale bar, 100 μm.

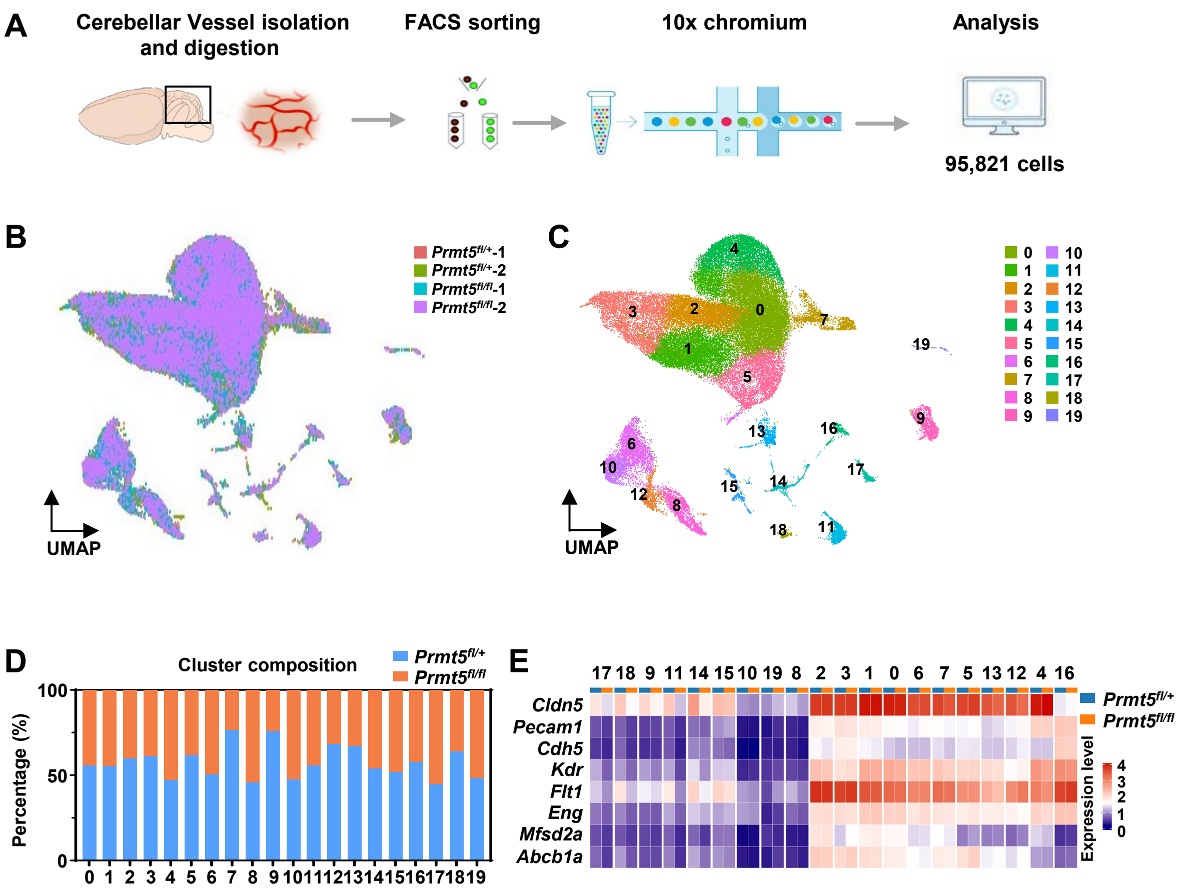


**Fig. S6. scRNA-sequencing of cerebellar ECs from *Prmt5*$^{fl/+}$ and *Prmt5*$^{fl/fl}$ mice**

(A) Schematic of the droplet-based 10x Genomics Chromium single-cell workflow.

(B) UMAP clustering colored according to group (24,209 *Prmt5*$^{fl/fl}$ cells and 31,911 *Prmt5*$^{fl/+}$ cells).

(C) UMAP plot showing identified cell subpopulations in an integrated analysis of *Prmt5*$^{fl/+}$ and *Prmt5*$^{fl/fl}$ groups.

(D) Graph illustrating the percentage distribution of *Prmt5*$^{fl/+}$ and *Prmt5*$^{fl/fl}$ cells within each cluster.

(E) Heatmap representing the expression of selected EC markers in *Prmt5*$^{fl/+}$ and *Prmt5*$^{fl/fl}$ groups across each cluster.

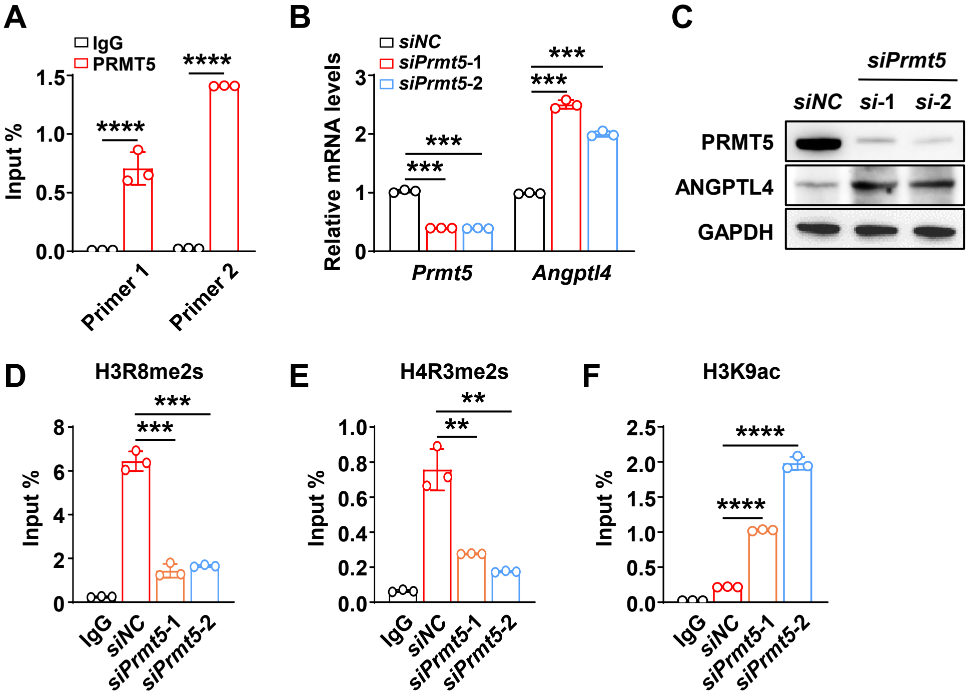


**Fig. S7. *Prmt5* negatively regulates *Angptl4* expression**

(A) ChIP-qPCR analysis of PRMT5 occupancy on the *Angptl4* promoter region in NIH-3T3 cells. ****P < 0.0001 (mean ± SEM, n = 3 per group).

(B) qPCR analysis of *Prmt5* and *Angptl4* mRNA levels in NIH-3T3 cells transfected with *siNC* or *siPrmt5*. ***P < 0.001 (mean ± SEM, n = 3 per group).

(C) Western blot analysis of PRMT5 and ANGPTL4 protein levels in NIH-3T3 cells transfected with *siNC* or *siPrmt5*. GAPDH served as the loading control.

(D-F) ChIP-qPCR analysis of the enrichment of H3R8me2s (D), H4R3me2s (E) and H3K9ac (F) at the *Angptl4* promoter region in NIH-3T3 cells transfected with *siNC* or *siPrmt5*. **P < 0.01, ***P < 0.001, ****P < 0.0001 (mean ± SEM, n = 3 per group).

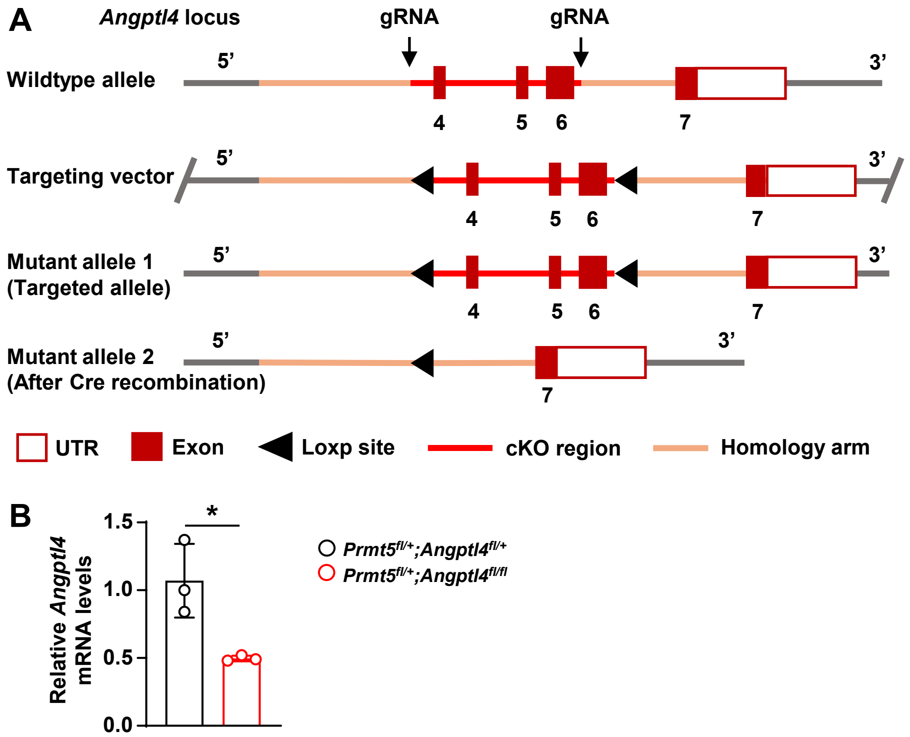


**Fig. S8. Generation of *Angptl4*$^{flox/flox}$ mice**

(A) Schematic diagram showing the strategy for generating *Angptl4*$^{flox/flox}$ mice.

(B) qPCR analysis of *Angptl4* expression in brain ECs isolated from P20 *Prmt5*$^{fl/+}$;*Angptl4*$^{fl/+}$ and *Prmt5*$^{fl/+}$;*Angptl4*$^{fl/fl}$ mice. *P < 0.05 (mean ± SEM, n = 3 per group).
